# Isoquinoline Alkaloids from Traditional Medicinal Plants of the Papaveraceae as a Third Line of Defense Against Bacterial Ocular Infections

**DOI:** 10.1101/2025.11.21.689706

**Authors:** Monika Dzięgielewska, Sylwia Zielińska, Bartłomiej Dudek, Malwina Brożyna, Weronika Kozłowska, Adam Matkowski, Marzenna Bartoszewicz, Yanfang Sun, Adam Junka

**Affiliations:** Dziegielewska Eye Institute, ul. Prymasa Augusta Hlonda 10c/u7, 02-972 Warsaw, Poland; ”P.U.M.A.” Platform for Unique Model Application, Department of Translative Technologies, Faculty of Pharmacy, Wroclaw Medical University, Borowska 211, 50-534 Wroclaw, Poland; Division of Pharmaceutical Biotechnology, Department of Pharmaceutical Biology and Biotechnology, Wroclaw Medical University, Borowska 211, 50-556 Wrocław, Poland; Division of Pharmaceutical Biology and Botany, Department of Pharmaceutical Biology and Biotechnology, Wroclaw Medical University, Borowska 211a, 50-556 Wrocław, Poland; Laboratory of Experimental Cultivation, Botanical Garden of Medicinal Plants, Wroclaw Medical University, Al. Jana Kochanowskiego 14, 50-556 Wrocław, Poland; Department of Pharmaceutical Microbiology and Parasitology, Wroclaw Medical University, Borowska 211a, 50-534 Wroclaw; Zhejiang International Joint Laboratory of Traditional Medicine and Big Health Products Development, Hangzhou, China; College of Life Sciences and Medicine, Zhejiang Sci-Tech University, Hangzhou, China

**Keywords:** *Papaveraceae* alkaloids, ocular infections, antibacterial activity, toxicity, translational phytotherapy

## Abstract

There is an evident need for novel ophthalmic therapeutics that operate outside the conventional antibiotic and chemically synthetized antiseptic paradigms. Medicinal plants represent a chemically rich and cost-effective source of bioactive compounds, though their use requires careful toxicological assessment. In this study, extracts from four *Papaveraceae* species - *Chelidonium majus* (roots), *Corydalis cheilanthifolia* (aerial parts and roots), *Glaucium flavum* (aerial parts and roots), and *Fumaria officinalis* (herb) - were investigated across three experimental levels: *in silico*, *in vitro*, and *in vivo*, using the *Galleria mellonella* larval model. Chemical profiling identified several isoquinoline alkaloids, including berberine, coptisine, protopine, chelidonine, allocryptopine, glaucine, and tetrahydropalmatine. *In silico* assessment suggested low systemic toxicity but possible blood-brain barrier permeability for some compounds. *In vitro* antimicrobial testing against *Staphylococcus aureus* ATCC 6538 and *Pseudomonas aeruginosa* ATCC 15442, performed using broth microdilution and bacterial-cellulose-based diffusion assays, demonstrated moderate but reproducible activity, particularly for *Ch. majus* and *C. cheilanthifolia* extracts. Cytotoxicity testing on fibroblast cell lines showed no significant toxicity in applied concentrations. *In vivo* testing showed that all extracts, except *C. cheilanthifolia* (herb_1) and *Ch. majus* (root) were virtually non-lethal to larvae, indicating a favorable safety profile. To evaluate their suitability for ocular application, wettability tests were performed using commercial tear substitute formulations. The incorporation of the *C*. *cheilanthifolia*_herb2 extract at bactericidal concentrations did not alter these physicochemical properties. Overall, the *Papaveraceae* extracts demonstrated moderate antibacterial activity and low cytotoxicity in an ophthalmologically relevant concentrations, supporting their potential use as adjuvant components in eye drop formulations - enhancing therapeutic efficacy, reducing required antiseptic dosage, and potentially lowering the risk of resistance development.

## Introduction

Antimicrobial resistance (AMR) has emerged as one of the most pressing global health threats of the 21st century. According to the landmark *Lancet* report prepared by the Global Burden of Disease AMR Collaborators, drug-resistant infections were directly responsible for an estimated 1.27 million deaths in 2019 and contributed to nearly 5 million more, surpassing the mortality burden of HIV/AIDS or malaria in the same period (Naghavi et al., 2024). This escalating crisis is driven by the remarkable genetic plasticity of bacteria, which enables the rapid acquisition of resistance determinants *via* horizontal gene transfer, genomic rearrangements, and mutational events. Selective pressure resulting from the excessive and often inappropriate use of antibiotics further accelerates the emergence of multidrug-resistant pathogens. In parallel with the erosion of antibiotic efficacy, antiseptics of synthetic origin have long been considered a robust alternative for topical infection control. Agents such as povidone-iodine, chlorhexidine, octenidine, or polyhexanide are widely used in clinical practice due to their broad activity spectrum, rapid bactericidal action, and initially presumed low propensity for resistance development. However, mounting evidence indicates that microorganisms exposed to repeated or sublethal doses of these agents may gradually acquire reduced susceptibility (Al-Doori et al., n.d.; Hardy et al., 2018; Kim et al., 2018). Reports of *Staphylococcus aureus* or *Pseudomonas aeruginosa* isolates exhibiting decreased responsiveness to chlorhexidine highlight that selective pressure is not limited to antibiotics alone (Hassan et al., 2013; Cieplik et al., 2019). Although the mechanisms underlying this phenomenon are multifactorial, ranging from efflux pump activation to alterations in cell envelope composition, the clinical consequence remains the same: antiseptics, once considered resistance-proof, are in the process of loosening their effectiveness. This underscores the urgent need to extend antimicrobial discovery efforts beyond conventional chemotherapeutics. These limitations necessitate the development of a “third line of defense” against resistant microorganisms. Beyond antibiotics and classical antiseptics (the first and the second line of defense, respectively) promising candidates include antimicrobial peptides of diverse origins and bioactive plant-derived products. Antimicrobial peptides represent an evolutionary defense strategy found in virtually all kingdoms of life, characterized by rapid, non-specific disruption of microbial membranes and additional immunomodulatory effects (Chakraborty et al., 2022). Likewise, plants produce a vast and chemically diverse arsenal of specialized metabolites, such as alkaloids, terpenoids, polyphenols (e.g flavonoids, tannins, stilbenoids) and other phytochemical classes - shaped by millions of years of co-evolution with microbial allies (microbiota) and antagonists (pathogens) (Anjali et al., 2023). This coevolution-driven “arms race” yields molecules that frequently act on microbial communication (quorum sensing) and pathogenicity (for example by blocking acyl-homoserine lactone signaling, down-regulating virulence genes, or disrupting biofilm architecture) rather than solely killing cells, which can lower the selective pressure for resistance. Well-documented examples include ajoene from garlic, which inhibits quorum-sensing regulated genes and biofilm formation, a broad family of phenolic compounds and flavonoids (e.g., quercetin, chlorogenic acid) that interfere with AHL-dependent signaling and biofilm development, and terpene-rich essential oil constituents (such as linalool and related terpenoids) that compromise biofilm integrity and bacterial membrane function (Nakamoto et al., 2020; Deryabin et al., 2019; Aditi et al., 2020; Saini et al., 2022). These anti-virulence and antibiofilm activities, reviewed extensively in the recent literature, make plant metabolites attractive candidates for sustainable, biocompatible adjuvants to conventional antimicrobials. At the same time, translation to human therapeutics requires rigorous toxicology, local pharmacokinetic profiling (absorption, retention and metabolism at the application site), and formulation work to address issues such as stability and poor oral/ocular bioavailability (Ezrari et al., 2025).

In this context, the eye provides a clinically relevant and highly sensitive model for exploring novel antimicrobial strategies within framework of “the third line of defense”. Ocular infections such as conjunctivitis, keratitis, blepharitis, and endophthalmitis are frequently associated with two major opportunistic pathogens: *S. aureus* and *P. aeruginosa*. The first of bacteria mentioned - a Gram-positive coccus - is notorious for its dual role as a commensal colonizer and an aggressive pathogen, with methicillin-resistant strains (MRSA) increasingly implicated in purulent conjunctivitis and postoperative infections. *P. aeruginosa*, in turn, is a Gram-negative bacillus characterized by extreme metabolic adaptability, rapid development of multidrug resistance, and a strong predilection for colonizing abiotic surfaces such as contact lenses and intraocular implants (Willcox, 2011; Teweldemedhin et al., 2017; O’Callaghan, 2018). Both species exhibit a remarkable ability to form biofilms - structured, extracellular-matrix-embedded microbial communities that confer enhanced tolerance to antibiotics, antiseptics, and host immune defenses (Uruén et al., 2020). Clinically, biofilm formation underlies the persistence and recurrence of ocular infections, complicates eradication efforts, and substantially increases the risk of vision-threatening sequelae. As such, ocular pathogens and their biofilm-associated phenotypes represent a stringent testing ground for evaluating the efficacy, safety, and translational potential of plant-derived antimicrobials.

Any attempt to utilize plant-derived antimicrobials as ophthalmic medication must be accompanied by careful toxicological scrutiny. The ocular surface is composed of highly specialized, delicate epithelial and stromal cells whose integrity is essential for maintaining vision. Even low cytotoxicity can result in irritation, impaired tear film stability, or delayed epithelial regeneration, while more pronounced toxicity may lead to corneal ulceration or permanent visual impairment. Unlike the skin or mucosa, ocular tissues are uniquely vulnerable due to their continuous exposure, high innervation, and limited regenerative capacity. Consequently, compounds with promising antimicrobial activity must be tested in parallel for their compatibility with human corneal and conjunctival epithelial cells, fibroblasts, and keratocytes (Müller and Kramer, 2008). Furthermore, ocular pharmacokinetics differ substantially from systemic or dermal exposure: absorption is modulated by tear turnover, blinking, and the barrier properties of the corneal epithelium. Therefore, safety assessment cannot rely solely on systemic toxicological data, but must specifically address local tolerability and bioavailability in the ophthalmic setting.

Formulation represents a critical step in implementation of plant-derived products into ophthalmic therapy. Topical eye drops remain the most widely used route of administration in ophthalmology, yet they suffer from rapid clearance through tear turnover and nasolacrimal drainage, resulting in short residence times and limited bioavailability. To achieve therapeutic efficacy, formulations must therefore balance physicochemical stability with favorable rheological properties that prolong ocular surface retention without compromising patient comfort or vision clarity. Approaches such as viscosity enhancers, mucoadhesive polymers, and in situ gelling systems have been extensively studied to optimize residence time. In this context, bacterial nanocellulose (BNC) polymer emerges as a promising next-generation carrier: it is biocompatible, highly hydrated, mechanically stable, and can serve as a depot for sustained release of hydrophilic and hydrophobic phytochemicals alike (Campano et al., 2025). Its transparent, non-irritating nature makes it ideally suited for ocular applications, where optical clarity and biocompatibility are essential. Embedding plant extracts into BNC-based formulations could enable controlled release and sustained antimicrobial activity, while paving the way for multifunctional ophthalmic preparations that integrate biofilm inhibition with anti-inflammatory and antioxidant effects.

Hence, plant natural products emerge as rational candidates for integration into ophthalmic formulations as most of the conventional antibiotics and synthetic antiseptics face escalating limitations. Some medicinal plants of the *Papaveraceae* family produce considerable amounts of isoquinoline alkaloids (IQA) with documented antimicrobial and anti-inflammatory potential (Zhang et al., 2024). *Corydalis cheilanthifolia* Hemsl., traditionally used in Asian phytotherapy for pain relief and wound healing, contains protoberberine and aporphine alkaloids such as corydaline and cheilanthifoline. Preliminary studies indicate antibacterial and antioxidant activity, although data on ocular applications remain scarce, whereas the reported toxicity appears generally low, with only sporadic suggestions of hepatotoxicity in other *Corydalis* species (Engman et al., 2023). *Chelidonium majus* L. (greater celandine), was reported since antiquity to cure blindness and other eye ailments and has been widely used in European folk medicine for the treatment of skin diseases, warts, and liver complaints (Zielinska et al. 2018). Its distinctive alkaloid profile includes chelidonine, sanguinarine, and chelerythrine and several other IQAs. These compounds exhibit strong antimicrobial and antibiofilm properties but the need for careful safety assessment in ocular contexts using credible approach is necessary (Greater Celandine, 2022; Li et al., 2024). *Fumaria officinalis* L., traditionally used in gastrointestinal and dermatological disorders, is rich in protopine and other IQA typical for the *Fumarioideae* subfamily and demonstrates spasmolytic, cholagogue, and antimicrobial effects. Toxicity is generally considered low, although hepatobiliary interactions cannot be excluded, and ophthalmic safety data remain limited (Ahmoda et al., 2025). *Glaucium flavum* Crantz, has been used in Mediterranean folk medicine for its expectorant and antitussive effects. Glaucine and related aporphine alkaloids from this plant display antimicrobial, bronchodilatory, and anti-inflammatory properties. At the same time, glaucine has been reported to cause central nervous system effects at higher systemic doses, and its local tolerability on epithelial cells has not been systematically evaluated (Akaberi et al., 2021). Collectively, these species represent rational candidates for exploratory ophthalmic formulations, combining evolutionary antimicrobial defenses with diverse phytochemical repertoires, yet their established or suspected toxicities emphasize the necessity of rigorous cytotoxicity testing on ocular cell lines prior to translational application.

In light of these considerations, the present study was designed to evaluate the potential of IQA from Papavearaceae plants extracts *in vitro* and *in vivo* as for a third line of defense agents against ocular pathogens, resistant to conventional therapies. In this work, extracts obtained from *C. cheilanthifolia*, *Ch. majus*, *F. officinalis*, and *G. flavum* were subjected to chromatographic profiling (HPLC) to characterize their chemical composition, followed by *in silico* toxicological analysis and microbiological testing against *S. aureus* and *P. aeruginosa* in both planktonic and biofilm forms. In parallel, cytotoxicity assays *in vitro* and *in vivo* were performed to ensure safety and tolerability, while formulation-oriented analyses explored the feasibility of incorporating these compounds into eye drop vehicles, specifically, bacterial nanocellulose-based systems. By integrating chemical, microbiological, toxicological, and formulation perspectives, this study aims to provide an attempt for assessing the translational potential of these traditional medicinal plants in ophthalmology.

## Material and Methods

### 1. Plant material

Plants were collected as shown in Table 1, dried at 35°C (Counter Inteligence, D-CUBE, LD-9013A dehydrator), and stored in airtight containers at room temperature till the extraction.

**Table 1.** Origin of Papaveraceae plant material used in the present study. ‘BGMP’–Botanical Garden of Medicinal Plants at the Wrocław Medical University, ‘Wilamowa’ – an experimental garden located in Wilamowa, Paczkow community, Nysa County, Poland. Herb – aerial parts containing green stems, leaves and flowers according to the pharmacopoeia guidelines.

| No. | Species | Part | Plant | GPS |
| --- | --- | --- | --- | --- |
| 1 | *Chelidonium majus* | Herb, Root | BGMP | 51.11737N<br>17.07483E |
| 2 | *Corydalis cheilanthifolia* | Herb (H1), Root | Wilamowa | 50.47449N<br>17.07382E |
| 3 | *Corydalis cheilanthifolia* | Herb (H2) | BGMP | 51.11737N<br>17.07483E |
| 4 | *Glaucium flavum* | Herb, Root | BGMP | 51.11737N<br>17.07483E |
| 5 | *Fumaria officinalis* | Herb | Wrocław, Niskie Łąki, spontaneous ruderal habitat | 51.09968N<br>17.06058E |

The voucher specimens were deposited in the herbarium of the Botanical Garden of Medicinal Plants of the Wroclaw Medical University under the codes:

> *Ch.m*._R_1–10/2023, *Ch.m*._H_1–10/2019, *C.ch*._R_1–10/2021, *C.ch*._H_1–10/2021, *C.ch*._H_1–10/2022, *G.f.*_R_1-3/2023, *G.f.*_H_1-3/2023, *F.o*._H_1-3/2023.

As a comparison plant material, a commercial sample of *Euphrasiae herba* – a traditional herbal drug used in Europe against eye complaints was purchased from Kawon-Hurt SJ (Authorization no. 2247, Gostyń, Poland)

### 2. Extraction and HPLC analysis

Plant material was crushed and powdered using a mortar and pestle and extracted with the mixture of MeOH (Stanlab, Poland) and 0.1% formic acid (Supelco, Germany) in a ratio 4:1 (v/v). Samples were extracted with ten parts of solvent per plant part. Half the volume of solvent was added to the plant material, sonicated (InterSonic, Poland) for 30 min at 25°C, and filtered through filter paper. The remaining solvent was added to the residual sediment, left for 24h at 4°C, sonicated (30 min), and filtered (Chempur, Poland). The extracts were combined and dried on a rotary evaporator (Buchi, Switzerland). The dried residue was dissolved in 10 ml of MeOH, transferred to test tubes and evaporated under the stream of nitrogen / lyophilized, and stored at 4°C until analysis. Dry extract was dissolved in DMSO (Sigma, Germany) in a concentration 1 mg/ml and further diluted with extraction solvent in a ratio 1:1. The High-Performance Liquid Chromatography with Diode-Array Detection (HPLC-DAD) analysis was performed using Agilent Technologies 1260 Infinity II chromatographic system (Agilent Technologies, Palo Alto, CA) coupled with Infinity War diode array detector. The separation was carried out using Kinetex C18 column (2.1 mm × 100 mm, 2.6µm, Phenomenex, United States). The mobile phase consisted of A: 10mM water solution of ammonium acetate (Fischer Chemical, Belgium) adjusted to pH 4 with glacial acetic acid ((Fischer Chemical, Belgium) and B: ACN (Merck, Germany). The elution program was as follows: 0-8 min 15-22%B; 8-18 min 22%B; 18-25 min 22-30%B; 25-28 min 30%B. Flow rate was 0,35 ml/min and the column was maintained in 25°C (±1°C). Detection was set at 280 nm. Identification of compounds was based on the comparison of the actual retention time of authentic reference standards (Chelidonine (later abbreviated as “CHD”), sanguinarine (SNG), berberine (BER), chelerithrine (CHL), protopine (PRO) – Extrasynthese, France; Allocryptopine (ACP) – Sigma-Aldrich, Germany; coptisine (COP) - PhytoLab GmbH, Germany). The data were acquired and processed using Agilent OpenLab Chemstation ver. 2.19.20. The references and data used for the identification of the investigated compounds is given in Supplementary Data 1.

### 3. The following bacterial strains were applied in this experiment

*S. aureus* Methicilin Resistant 33591 and *P. aeruginosa* 15542 from American Type Culture Collection, (ATCC, Manassas, VA, USA), *Komagataeibacter xylinus* ATCC 53524 (American Type Culture Collection, Manassas, VA, USA)

### 4. *In silico* ADME profiling of alkaloids detected by HPLC

The pharmacokinetic properties of alkaloids identified in plant extracts were evaluated using the SwissADME web tool (http://www.swissadme.ch Swiss Institute of Bioinformatics, Lausanne, Switzerland). For each compound, the canonical SMILES notation was retrieved from the PubChem database (National Center for Biotechnology Information, Bethesda, MD, USA) and introduced as input. SwissADME provided predictions of physicochemical descriptors (molecular weight, number of rotatable bonds, topological polar surface area [TPSA], hydrogen bond donors and acceptors), lipophilicity (consensus logP), water solubility, and drug-likeness filters (Lipinski, Ghose, Veber, Egan, and Muegge rules). Absorption, distribution, metabolism, and excretion (ADME) parameters were evaluated, including gastrointestinal (GI) absorption, blood–brain barrier (BBB) permeability, and substrate/inhibitor status for P-glycoprotein (P-gp) and major cytochrome P450 (CYP) isoforms (CYP1A2, CYP2C19, CYP2C9, CYP2D6, and CYP3A4). In addition, the BOILED-Egg model implemented in SwissADME was applied to visualize passive gastrointestinal absorption and BBB penetration. Predicted pharmacokinetic profiles were exported and summarized for all alkaloids, enabling comparative evaluation of their oral bioavailability and potential for systemic distribution.

### 5. *In silico* prediction of alkaloids toxicity

The toxicological profiles of selected alkaloids identified in *Papaveraceae* extracts were assessed using the ProTox-III webserver (https://tox.charite.de/protox3/) Charité – Universitätsmedizin Berlin, Germany). Canonical SMILES notations of each alkaloid were retrieved from the PubChem database (National Center for Biotechnology Information, Bethesda, MD, USA) and submitted as input. For each molecule, ProTox-II provided predicted values of acute oral toxicity (LD₅₀ in mg/kg body weight) and toxicity class according to the Globally Harmonized System (GHS) of classification. In addition, endpoints covering hepatotoxicity, cytotoxicity, carcinogenicity, mutagenicity, immunotoxicity, and Tox21 pathway activation (nuclear receptor signaling and stress response pathways) were generated. Probabilities of compound assignment to toxic versus non-toxic categories were reported as confidence scores. The resulting predictions were compiled for all tested alkaloids. This approach allowed a rapid, systematic estimation of the toxicological risk profile of alkaloids prior to further *in vitro* and *in vivo* validation.

### 6. Estimation of Minimal Inhibitory Concentration of plant extracts using microdilution method

Dried plant extracts were dissolved in DMSO (analytical grade, Sigma-Aldritch, Germany) to obtain 5mg/mL stock solutions, which were subsequently diluted twice with sterile deionized water to working intermediates and finally subjected to two-fold serial dilution in cation-adjusted Mueller–Hinton broth (CA-MHB). The final DMSO content in assay wells did not exceed 1% (v/v), and solvent controls were included on every plate. Bacteria were grown overnight on TSA, suspended in saline, and adjusted to 0.5 McFarland using a densitometer (DEN-1B, SIA Biosan, Latvia), followed by dilution into CA-MHB to yield a final inoculum of approximately 5 × 10^5 CFU/mL per well. Minimum inhibitory concentrations (MICs) were determined in 96-well flat-bottom plates (Wuxi Nest Biotechnology, China) by the standard broth microdilution method. Plates were inoculated and incubated at 37 °C for 24 h. Following incubation, 2,3,5-triphenyl-tetrazolium chloride (TTC; 0.05–0.10% w/v, Chempur, Poland) was added to each well, and plates were further incubated for 30 min. The MIC was defined as the lowest extract concentration at which no red coloration developed, indicating the absence of TTC-formazan formation. The result was verified by spot method. Each assay plate included growth, sterility, and solvent controls. All experiments were carried out in three independent biological replicates on separate days, with 3 technical duplicates included in each run. MIC values were recorded as the modal outcome across repeats; in cases where values differed by one dilution step, the higher MIC was reported.

### 7. Estimation of Minimal Biofilm Eradication Concentration of plant extracts using the microdilution method

Overnight cultures grown on tryptic soy agar were suspended in saline and adjusted to 0.5 McFarland (DEN-1B, SIA Biosan, Latvia), then diluted into CA-MHB to obtain ∼5 × 10^5 CFU/mL. Biofilms were established in 96-well flat-bottom polystyrene plates (Wuxi Nest Biotechnology, China) by inoculating 100 µL per well and incubating at 37°C for 24 h without agitation. After incubation, medium was carefully aspirated, wells were gently rinsed once with sterile PBS to remove non-adherent cells, and 200 µL of fresh CA-MHB containing the extract dilution series (or controls) was added to each well. Plates were incubated a further 24 h at 37 °C. Following treatment,; 0.05–0.10% TTC w/v) was added to each well and plates were incubated for 30 min. The MBEC was defined as the lowest extract concentration at which no red coloration developed, indicating absence of TTC-formazan formation by the biofilm. In parallel, eradication was corroborated by viable-count “spot” readout following biofilm disruption (PBS rinse, mechanical scraping of the well contents, then spotting on Mueller–Hinton agar). Each plate included growth, sterility, and solvent controls. Experiments were performed as three independent biological replicates on separate days, each with 3 technical duplicates. When replicate MBECs differed by one two-fold dilution, the higher value was reported.

### 8. Preparation of Bacterial Nano-Cellulose carrier containing the plant extracts

Bacterial nanocellulose (BNC) discs were produced by *Komagataeibacter xylinus* ATCC 53524 (American Type Culture Collection, Manassas, VA, USA) cultured in Hestrin–Schramm (HS) medium under static conditions at 28 °C for 7 days. Following incubation, 11 mm in diameter pellicles were harvested, and purified by immersion in 0.1 M NaOH (Chempur, Poland) at 80 °C until all bacterial cells and debris were removed. The discs were then thoroughly rinsed with sterile water until neutral pH was achieved. Sterilization was performed by autoclaving at 121 °C for 15 min. Prior to use in antimicrobial assays, BNC discs were equilibrated in sterile deionized water and stored at 4 °C under aseptic conditions. For disk diffusion testing, sterile discs were transferred into wells of sterile 24-well plates and soaked with plant extract solutions for 24 h at room temperature, during which each disc absorbed on average 600 µL of extract solution, displacing the water naturally present in the hydrated matrix.

### 9. Evaluation of Antibacterial Activity of extracts using Modified Disk Diffusion Assay

Antimicrobial activity of the extracts was assessed using a modified disk diffusion assay in which BNC discs replaced conventional paper carriers. BNC discs with a diameter matching the wells of 24-well plates (≈11 mm) were first sterilized and then saturated with plant extracts by incubation for 24 h in extract solutions prepared from DMSO stocks diluted in sterile deionized water (final DMSO ≤1% v/v). During the soaking process, each disc absorbed on average 600 µL of solution, which effectively displaced the water naturally present in the hydrated BNC matrix. After impregnation, discs were gently blotted under sterile conditions to remove surface excess liquid and immediately transferred onto Mueller–Hinton agar plates (Biomaxima, Poland) previously inoculated with standardized bacterial suspensions. The inoculum was prepared from overnight cultures of *S. aureus* ATCC 6538 and *P. aeruginosa* ATCC 15442 (American Type Culture Collection, Manassas, VA, USA), adjusted to 0.5 McFarland with a DEN-1B densitometer (SIA Biosan, Latvia), and uniformly spread onto the agar surface. Plates with BNC discs were incubated at 37 °C for 18–20 h. After incubation, the inhibition zones were photographed under standardized lighting conditions and analyzed using ImageJ software (NIH, USA). The measurement scale was calibrated for each image using the “Set Scale” function based on a millimeter reference. The total area of bacterial cellulose (BC) discs was subtracted from the measured inhibition halo to quantify only the zone of microbial growth inhibition. Final results were expressed as inhibition area (mm²). Each plate included solvent controls (BC discs soaked in DMSO at the highest applied concentration), sterility controls, and reference antimicrobials. All experiments were performed in two independent biological replicates, each in three technical duplicates, and data were presented as mean inhibition area ± standard deviation.

### 10. Interaction modeling (Bliss independence and Highest Single Agent, HSA)

Interaction analyses were performed on inhibition magnitudes obtained in the modified BNC disk-diffusion assay for *S. aureus* and *P. aeruginosa*. For each extract, six independent replicates were photographed and quantified in ImageJ; the area corresponding to the 12-mm cellulose carrier was subtracted to yield the net inhibition area, which was used consistently across singles and combinations. Single-agent activities were taken directly where a single dominant alkaloid was present (PRO from *F. officinalis* herb, CHD from *Ch. majus* extract being collected in DoTT collection (Supplementary Data), glaucine, GLA from *G. flavum* herb) and, where not available, reconstructed within the same species and experimental run by algebraic subtraction (COP = *C. cheilanthifolia* H1 [COP+PRO+BER] − *C. cheilanthifolia* H2 [PRO+BER]; BER = *C. cheilanthifolia* H2 [PRO+BER] − PRO solo; THP = *G. flavum* root [PRO+CHD+THP+GLA] − (PRO solo + CHD solo + GLA solo)); when ACP was present without a reliable single, it was treated as unknown and omitted from model predictions as a conservative choice. Computations were carried out in a spreadsheet-based analytical tool implementing the closed-form equations described below: the largest observed inhibition magnitude within each species was taken as Emax; each single-agent magnitude Mi was converted to a fractional effect Ei = Mi / Emax; the Bliss reference for a combination C was EBliss(C) = 1 − ∏_{i∈C}(1 − Ei) and was converted back to the original unit as MBliss(C) = EBliss(C) × Emax; the HSA reference was MHSA(C) = max_{i∈C} Mi. For each combination we calculated ΔBliss = Mobs − MBliss and ΔHSA = Mobs − MHSA and assigned interaction classes with a symmetric tolerance of ±10% of the observed value: synergy if Δ > 0.10 × Mobs, additivity if |Δ| ≤ 0.10 × Mobs, antagonism if Δ < −0.10 × Mobs; “HSA vs Bliss” was reported as agreement when both models returned the same class and as disagreement otherwise. All intermediate calculations were maintained at ≥4 significant digits and rounded to two decimals in the reported tables.

### 11. Assessment of in vitro cytotoxicity of extract toward L929 cell using Neutral MTT test

Dulbecco’s modified Eagle’s medium (DMEM, Biowest, Riverside, MO, USA) supplemented with 1% (v/v) amphotericin B (Biowest, Riverside, MO, USA), 1% (v/v) penicillin with streptomycin (Biowest, Riverside, MO, USA), and 10% (v/v) fetal bovine serum FBS (Biowest, Riverside, MO, USA) was used for cells’ culturing. A suspension of the L929 murine fibroblast cell line (ATCC CCL) at a density of 1×10^5/mL was prepared, and 100 µL was added to the wells of 96-well plates (VWR, Radnor, PA, USA). The plates were incubated for 24 h/37°C, 5% CO2 (Binder C 150 UL CO2 incubator, Germany). Next, the medium was gently removed. The stock solutions of the compounds were geometrically diluted in medium at the concentration ranging from 0.05mg/mL to 0.002mg/mL and transferred to the cells at a volume of 100 µL. The following control samples were prepared: growth control and control of DMSO (dimethyl sulfoxide, Sigma Aldrich, Germany) geometrically diluted in medium at concentrations ranging from 1% (v/v) to 0.03% (v/v). The plates were retransferred to the incubator for 24 h. After that, the medium was removed and 100 µL of the solution of 1mg/mL MTT ((3-(4,5-dimethyl-2-thiazolyl)-2,5-diphenyl-2H-tetrazolium bromide) (Sigma Aldrich, Germany) in DMEM was added for 2h, and the plates were incubated under the same conditions. Subsequently, the dye was removed, and 100 µL DMSO was added for 30 min to dissolve crystals (shaking at 400 rpm (Plate Shaker-Thermostat PST-60HL-4 (Biosan, Latvia)). The absorbance of the solution was then measured at 570 and 630 nm with a spectrophotometer (Multiscan® GO, Thermo Scientific, USA). The values at 630 nm were subtracted from the values at 570, and the viability of the treated cells was calculated as follows: Cells Viability [%] = (AbS /Ab C) x 100%; AbS-mean absorbance value of treated samples; AbC- mean values of the growth control. The experiment was performed in three independent experiments, each with 4 repetitions.

### 12. Assessment of *in vivo* cytotoxicity using *Galleria mellonella* larval model

The *in vivo* efficacy and toxicity of the extracts were examined using larvae of *G. mellonella* reared in the P.U.M.A. laboratory (Wroclaw Medical University, Poland). The larvae originated from a standardized in-house culture maintained under controlled conditions (30°C, constant humidity, darkness) and fed a uniform artificial diet to ensure physiological consistency across experiments. Groups of six larvae per compound (weighing 250–300 mg each, free of melanization or physical damage) were injected into the last left proleg with 10 µL of extract solution using a Hamilton syringe (Hamilton, Bonaduz, Switzerland). Control groups received either phosphate-buffered saline or DMSO at the corresponding maximal concentration. A 75% ethanol solution was used as a positive (toxicity) control to verify test responsiveness. Larvae were incubated at 37 °C in the dark and monitored for survival every 24 h for up to 96 h.

### 13. Commercial Ophthalmic Preparations

Three commercially available eye drop formulations were included in the study. Dexoftal® (TEGE Pharma B.V, Breda, Netherlands), hereafter referred to as DX, contains dexpanthenol (2%) polihexanid 0.0001% and hydroxypropylcellulose (HydraFlex™, 0.5%) as viscosity enhancer and mucoadhesive polymer. Thealoz Duo® (Laboratoires Théa, Clermont-Ferrand, France), abbreviated as TD, is composed of trehalose (3%), sodium hyaluronate (0.15%), trometamol, sodium chloride, hydrochloric acid, and water for injection. Hialeye Free® (Adamed Pharma, Czosnow, Poland), denoted as HF, contains sodium hyaluronate (0.2%), sodium phosphate dodecahydrate, sodium dihydrogen phosphate monohydrate, sodium chloride, and water. All products were used within their shelf life and handled under sterile conditions.

### 14. The wettability of ophthalmic preparations

**The wettability** of the tested ophthalmic formulations (DX, TD, HF) was evaluated by contact angle analysis performed on surfaces with different physicochemical properties. Droplets of 40 μL were dispensed using a 100 μL microsyringe (Hamilton, Reno, NV, USA) onto three substrates: hydrophobic poly(L-lactic acid) (PLLA) plates (Goodfellow, Cambridge, UK), (ii) hydrophilic glass slides (Thermo Fisher Scientific, Waltham, MA, USA), and an *ex vivo* tear film model. The artificial tear film consisted of sodium chloride (6.8 g/L), potassium chloride (1.4 g/L), calcium chloride dihydrate (0.08 g/L), magnesium chloride hexahydrate (0.1 g/L), sodium bicarbonate (2.2 g/L), glucose (1 g/L), and mucin (0.5% w/v; porcine stomach type II, Sigma-Aldrich, St. Louis, MO, USA) dissolved in phosphate-buffered saline (PBS, pH 7.4). Measurements were performed at 24 °C under ambient laboratory conditions. For each formulation–substrate combination, at least six replicates were analyzed. Images of sessile drops were photographed and contact angles were calculated using ImageJ software (NIH, Bethesda, MD, USA) with the Contact Angle plug-in (code has been written by Marco Brugnara and itï¿½ï¿½s based on the plug-in "Pointpicker”; https://imagej.net/ij/plugins/contact-angle.html).

### 15. Combination of Plant Extracts with Chosen Commercial Eye Drop

To assess the feasibility of incorporating plant extracts into existing ophthalmic vehicles, *C. cheilanthofolia* H2 extract was combined with DX tear drop. The extract was added to the formulation to achieve a final concentration corresponding to 2 × MIC, as determined in the broth microdilution assay. The resulting mixture was used to saturate sterile BNC discs for 24 h, as described previously. For comparison, parallel BNC discs were prepared with the same extract diluted in 1% DMSO and water; another control was BNC saturated with 1% DMSO and water. Antimicrobial activity was subsequently evaluated using the modified disk diffusion method (Section Material and Methods 8).

### 16. Statistical Analysis

Statistical analyses were performed using GraphPad Prism 10 (San Diego, CA, USA). Normality of distribution was verified using Shapiro–Wilk’s test. An Analysis of Variance (ANOVA) was performed to assess statistical significance. For multiple comparisons, Tukey’s or Sedak’s post hoc test was applied. A p-value threshold of less than 0.05 was set for significance in the ANOVA. For the Tukey post hoc analysis, significance levels were further categorized as p < 0.001 and p < 0.0001 for specific pairwise comparisons.

## 3. Results

At the initial stage of the study, phytochemical profiling of extracts obtained from Papaveraceae species was performed in order to identify and quantify dominant alkaloids. HPLC analysis revealed the presence of composition of alkaloids depending on the plant species and the part of the plant (herb or root). The detected ISQ alkaloids included ACP, BER, GLA, COP, PRO, THP, as well as their derivatives. The concentrations of detected alkaloids are provided in **Table 2**.

**Table 2.** Identified alkaloids and their concentrations (µg/mg DW ± SD) in extracts of *Papaveraceae* species (herb and root fractions). **DW** -dry weight; **SD** - standard deviation. **H** – herb (aerial parts); **R**-root. H1 – origin Wilamowa, H2 – origin BGMP Wroclaw

| extract | alkaloid | average (µg/mg DW) ±SD; n=3 |
| --- | --- | --- |
| *C. cheilanthifolia*_H1 | Protopine | 8.24±0.08 |
|  | Coptisine | 4.14±0.02 |
|  | Protopine derivative | 8.11±0.32 |
|  | Berberine | 10.48±0.28 |
| *C. cheilanthifolia*_H2 | Protopine | 13.37±0.14 |
|  | Protopine derivative | 12.34±0.10 |
|  | Berberine | 7.32±0.15 |
| *C. cheilanthifolia*_R | Protopine | 35.92±0.22 |
|  | Coptisine | 11.56±0.11 |
|  | Protopine derivative | 11.18±0.20 |
| *Ch. majus*_R | Protopine | 44.84±0.61 |
|  | Allocriptopine | 5.62±0.09 |
|  | Chelidonine | 3.38±0.12 |
|  | Coptisine | 69.82±0.55 |
|  | Berberine | 5.45±0.02 |
| *F. officinalis*_H | Protopine | 4.12±0.07 |
| *G. flavum*_H | Glaucine derivative 1 | 3.15±0.06 |
|  | Glaucine derivative 2 | 10.03±1.16 |
|  | Glaucine | 170.04±20.36 |
| *G. flavum*_R | Glaucine derivative | 3.08±0.03 |
|  | Protopine | 61.16±0.23 |
|  | Chelidonine | 4.64±0.07 |
|  | Tetrahydropalmatine | 3.67±0.09 |

| predicted lethal dose (LD<sub>50</sub>, mg/kg)<br>[ProTox III] |  |  |  |  |  |  |
| --- | --- | --- | --- | --- | --- | --- |
| PRO | BER | CHD | GLA | COP | ACP | THP |
| 940 | 200 | 460 | 400 | 200 | 940 | 400 |

**Table 3.** Interaction analysis of alkaloid-rich extracts from *Papaveraceae* species against *S. aureus* based on Bliss independence and Highest Single Agent (HSA) models (±10%). The table summarizes experimental inhibition zones (mm²) and predicted activities calculated using Bliss and HSA models. Deviations (Δ) between observed and predicted values were used to classify the type of interaction: Synergy – observed value exceeds prediction by >10%; Additivity - within ±10%; Antagonism - observed value < prediction by >10%. “HSA vs Bliss” indicates whether both models provided consistent (**Ag** - agreement) or divergent (**D** - disagreement) classifications. Single-agent activities used for the analysis (in mm): protopine (**PRO**) - 6.735, chelidonine (**CHED**) - 19.479, glaucine (**GLA**) - 12.953, coptisine (**COP**) - 14.866, berberine (**BER**) - 16.004, tetrahydropalmatine (**THP**) - 15.427.

| Plant | Alkaloid | Observed Zone (mm) | Bliss Prediction (mm) | ΔBliss (Obs – Bliss) [mm] | Bliss Interpretation ($\pm 10\%$) | HSA Prediction (mm) | ΔHSA (Obs – HSA) [mm] | HSA Interpretation ($\pm 10\%$) | HSA vs Bliss |
| --- | --- | --- | --- | --- | --- | --- | --- | --- | --- |
| *C. cheilant hifolia* H1 | cop +pro +ber | 49.96 | 38.73 | 11.23 | SYN | 29.4 | 20.56 | SYN | **Ag** |
| *C. cheilant hifolia* H2 | pro +ber | 33.21 | 31.23 | 1.98 | ADD | 29.4 | 3.81 | SYN | **D** |
| *C. cheilant hifolia* R | cop +pro | 23.76 | 19.43 | 4.33 | SYN | 16.75 | 7.01 | SYN | **Ag** |
| *Ch. majus* R | cop +pro +ber +chel +acp | 35.05 | 40.03 | -4.98 | ANT | 29.4 | 5.65 | SYN | **D** |
| *F. officinalis* H | pro | 3.81 | 3.81 | 0.0 | ADD | 3.81 | 0.0 | ADD | **Ag** |
| *G. flavum* R | pro +chel +thp +gla | 56.53 | 41.2 | 15.33 | SYN | 30.35 | 26.18 | SYN | **Ag** |
| *G. flavum* H | gla | 18.22 | 18.22 | 0.0 | ADD | 18.22 | 0.0 | ADD | **Ag** |

**Table 3. Interaction analysis of alkaloid-rich extracts from *Papaveraceae* species against *P.**
| Plant | Alk | Observed Zone (mm) | Bliss Prediction (mm) | ΔBliss (Obs – Bliss) [mm] | Bliss Interpretation ($\pm 10\%$) | HSA Prediction (mm) | ΔHSA (Obs – HSA) [mm] | HSA Interpretation ($\pm 10\%$) | HSA vs Bliss |
| --- | --- | --- | --- | --- | --- | --- | --- | --- | --- |
| *C. cheilant hifolia* H1 | cop +pro +ber | 37.6 | 29.98 | 7.62 | **SYN** | 16.0 | 21.6 | **SYN** | **Ag** |
| *C. cheilant hifolia* H2 | pro +ber | 22.74 | 20.76 | 1.98 | **ADD** | 16.0 | 6.73 | **SYN** | **D** |
| *C. cheilant hifolia* R | cop +pro | 23.74 | 19.77 | 3.97 | **SYN** | 14.87 | 8.88 | **SYN** | **Ag** |
| *Ch. majus* R | cop +pro +ber +chel +acp | 38.05 | 38.76 | -0.71 | **ADD** | 19.48 | 18.57 | **SYN** | **D** |
| *Ch. majus* H | pro | 19.48 | 19.48 | -0.0 | **ADD** | 19.48 | 0.0 | **ADD** | **Ag** |
| *F. officinalis* H | pro +chel +thp +gla | 6.73 | 6.73 | 0.0 | **ADD** | 6.73 | 0.0 | **ADD** | **Ag** |
| *G. flavum* R | gla | 54.59 | 37.75 | 16.84 | **SYN** | 19.48 | 35.11 | **SYN** | **Ag** |
| *G. flavum* H | cop +pro +ber | 12.95 | 12.95 | 0.0 | **ADD** | 12.95 | 0.0 | **Additivity** | **Ag** |

Extracts from *C. cheilanthifolia* were characterized predominantly by protopine, berberine, and coptisine, with notably higher concentrations of protopine in the roots compared to the aerial parts. The extract of *Ch. majus* roots was dominated by coptisine and protopine, together with detectable allocryptopine, berberine and chelidonine. *F. officinalis* extract was chemically simpler, with protopine as the only major alkaloid identified. In contrast, *G. flavum* displayed the highest total alkaloid content, with glaucine exceeding 170 µg/mg DW in the aerial part, and high protopine content (over 60 µg/mg DW) in the root. Collectively, these results indicate that each *Papaveraceae* species exhibits a distinct alkaloid fingerprint, with *G. flavum* herb and *Ch. majus* root extracts representing the richest sources. Berberine and coptisine were assigned the lowest LD₅₀ values (200 mg/kg), corresponding to GHS (Globally Harmonized System) class 3, while chelidonine (460 mg/kg), glaucine and tetrahydropalmatine (both 400 mg/kg) showed intermediate toxicity (class 3). Protopine and allocryptopine were classified as less toxic, with predicted LD₅₀ values of 940 mg/kg (class 4).

Next, the SwissADME BOILED-Egg analysis was perform to check the properties of above-enlisted active compounds in human body and revealed clear differences in the predicted pharmacokinetic profiles of the tested alkaloids (**Figure 1**). Allocryptopine, chelidonine, and protopine were located within the white region, indicating high probability of gastrointestinal absorption but no blood–brain barrier (BBB) penetration. In contrast, glaucine and tetrahydropalmatine clustered within the yellow region, consistent with favorable BBB penetration and thus potential neuroactive properties. Berberine and coptisine were positioned outside both regions, suggesting poor oral absorption and negligible BBB access. Regarding interaction with P-glycoprotein (P-gp), compounds represented as blue dots are predicted to be P-gp substrates and therefore actively effluxed, whereas red dots denote non-substrates with higher likelihood of intracellular accumulation.

**Figure 1.**
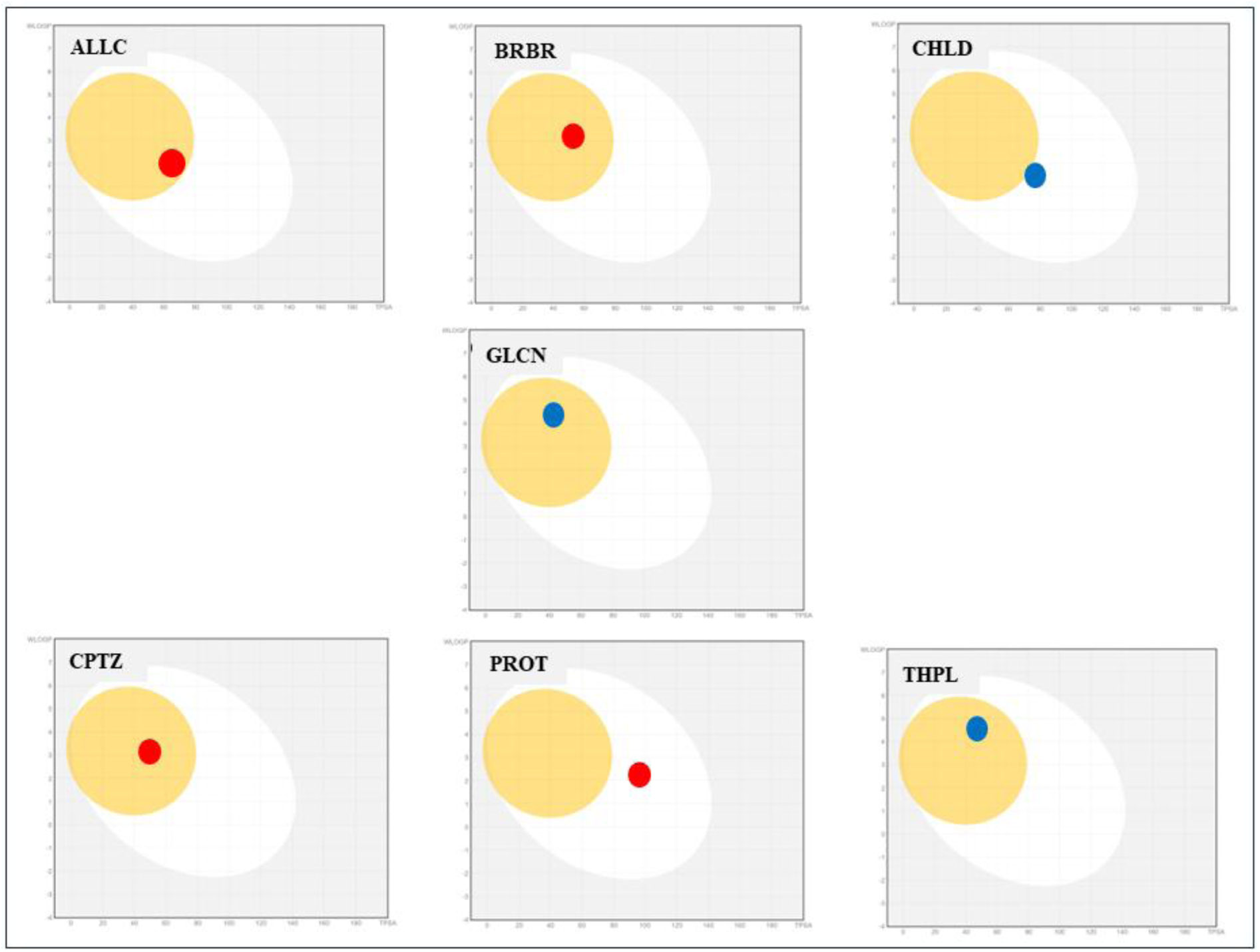
BOILED-Egg plots of *Papaveraceae* alkaloids generated with SwissADME. The position of each dot indicates the predicted pharmacokinetic behavior: location of dot within the yellow region (“yolk”) corresponds to a high probability of blood–brain barrier (BBB) permeability; within the white region (“egg white”) to a high probability of gastrointestinal absorption; and outside both regions to low probability of either property. Dot color indicates interaction with P-glycoprotein (P-gp): red dots represent compounds predicted not to be P-gp substrates, while blue dots represent compounds predicted to be P-gp substrates, i.e., subject to active efflux and limited central nervous system accumulation. Abbreviations: **ACP** – allocryptopine, **BER** – berberine, **CHD** – chelidonine, **GLA** – glaucine, **COP** – coptisine, **PRO** – protopine, **THP** – tetrahydropalmatine.

While the BOILED-Egg model provided insights into the pharmacokinetic behavior of the investigated alkaloids, pharmacological feasibility must be balanced with their toxicological safety. Therefore, in parallel with ADME profiling, we employed the ProTox-III platform to predict no only general toxicity (as presented in **Table 1**) but also organ-specific and systemic toxicities based on structural alerts (**Figure 2**, **Figure 3**).

**Figure 2.**
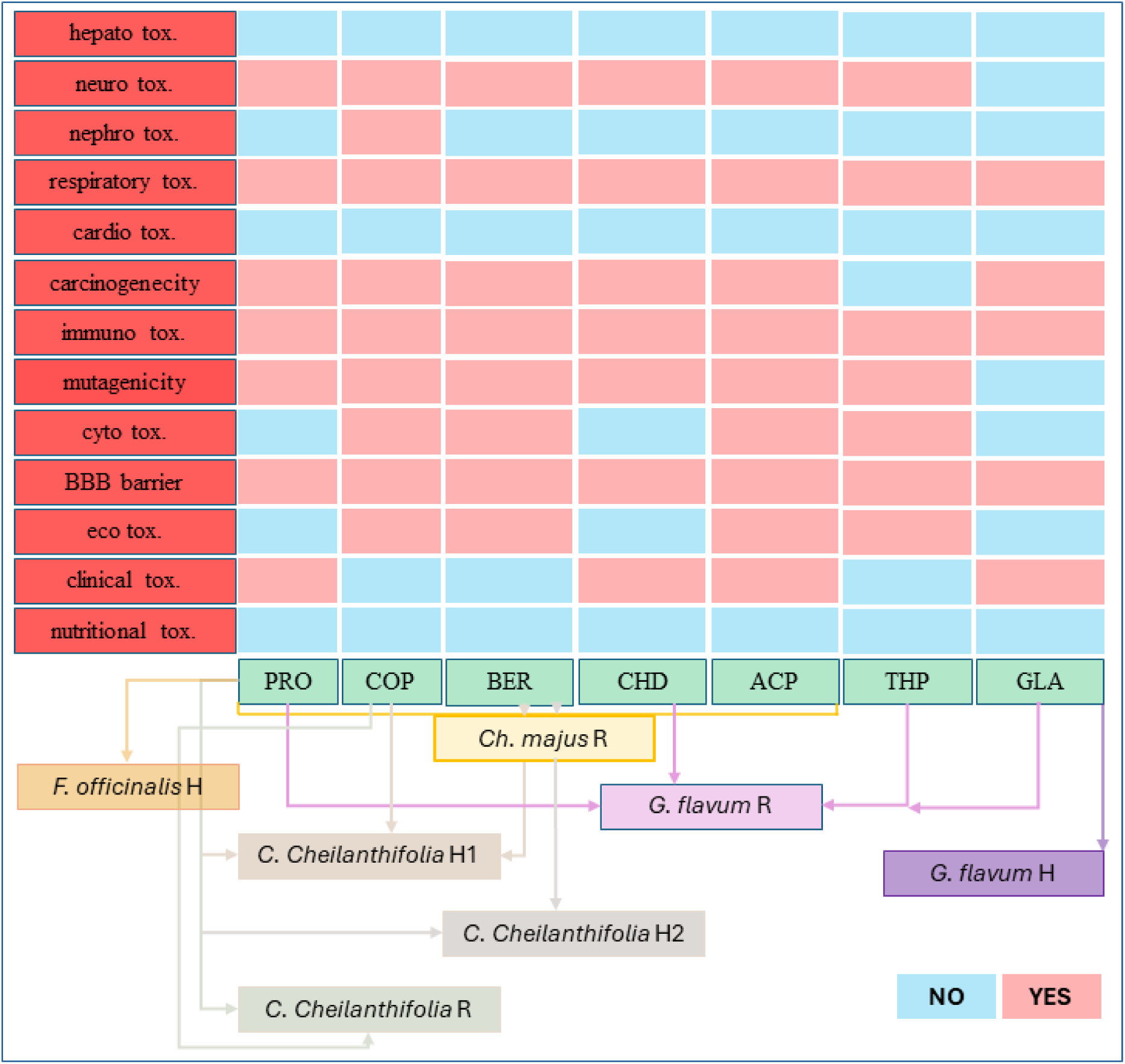
Predicted organ-specific and systemic toxicity profiles of major *Papaveraceae* alkaloids based on ProTox-III analysis. Heatmap summarizing the predicted toxicological endpoints for alkaloids detected in *Papaveraceae* extracts. Red fields indicate a positive (toxic) prediction, while blue fields denote absence of toxic activity. The lower part of the diagram links individual compounds to the plant species and plant parts in which they were identified. Abbreviations: **PRT** - protopine; **CPTZ** - coptisine; **BRBR** - berberine; **CHLD** - chelidonine; **ALLC** - allocryptopine; **THPL** - tetrahydropalmatine; **GLCN** - glaucine; **H** - herb; **R** - root. Toxicity endpoints: **hepato tox**.- hepatotoxicity; **neuro tox**. – neurotoxicity; **nephro tox.** - nephrotoxicity; **respiratory tox.** - respiratory toxicity; **cardio tox.** - cardiotoxicity; **carcinogenicity** - carcinogenic potential; **immuno tox.** - immunotoxicity; **mutagenicity** - genotoxic potential; **cyto tox**. - cytotoxicity; **BBB barrier** - blood–brain barrier permeability; **eco tox.** - ecological toxicity; **clinical tox.** - predicted clinical toxicity; **nutritional tox.** - nutritional imbalance-related toxicity.

**Figure 3.**
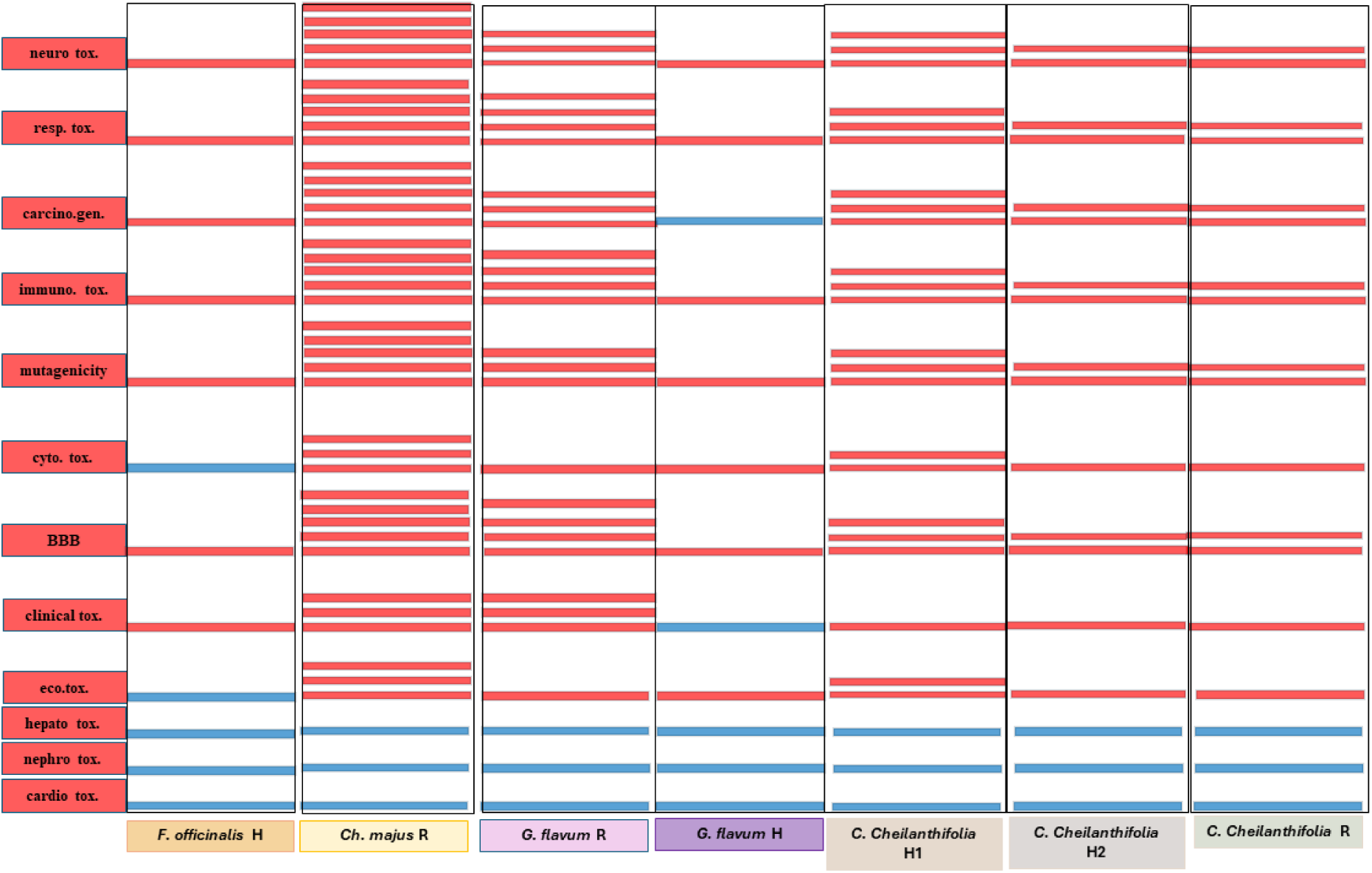
Cumulative *in silico* toxicity profiles of Papaveraceae extracts predicted using the ProTox-III platform. Each column corresponds to an extract obtained from *Fumaria officinalis* (H – herb), *Chelidonium majus* (H – herb, R – root), *Glaucium flavum* (H, R), and *Corydalis cheilanthifolia* (H1, H2, R). Each horizontal red bar represents one alkaloid identified within a given extract that was predicted to be active for a specific toxicity endpoint. The number of red bars therefore reflects the number of alkaloids exhibiting the given type of toxicity, whereas blue fields indicate that none of the alkaloids present in the extract were predicted as toxic in that category. Evaluated endpoints include **neuro tox.** – neurotoxicity; **resp. tox.** – respiratory toxicity; **carcino.gen.** – carcinogenicity; **immuno. tox.** – immunotoxicity; **mutagenicity** – genotoxic potential; **cyto. tox.** – cytotoxicity; **BBB** – blood–brain barrier permeability; **clinical tox.** – predicted systemic clinical toxicity; **eco. tox.** – ecological toxicity; **hepato tox.** – hepatotoxicity; **nephro tox.** – nephrotoxicity; and **cardio tox.** – cardiotoxicity.

**Figure 4.**
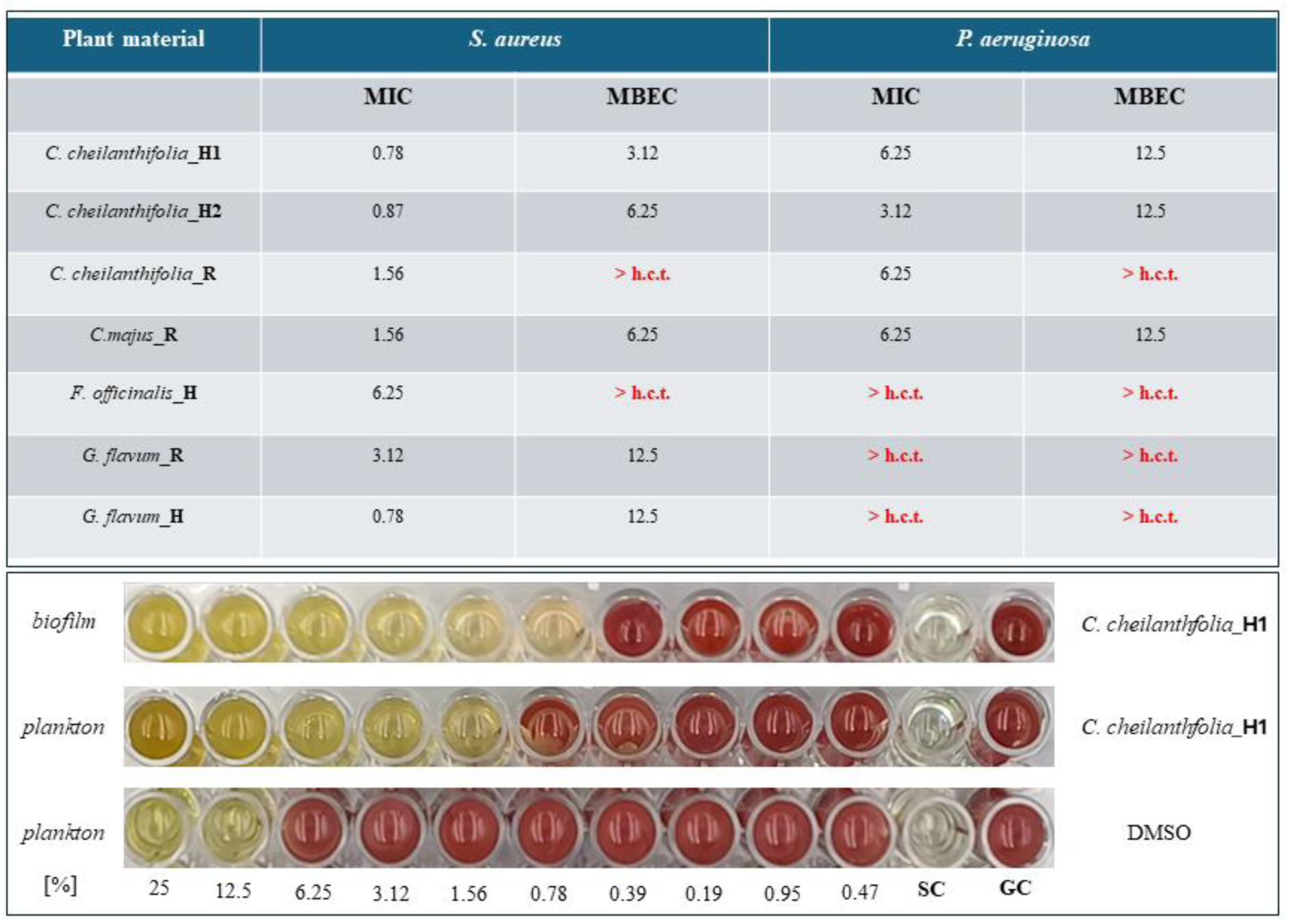
Antimicrobial activity of plant extracts against *S. aureus* and *P. aeruginosa*. The table shows minimum inhibitory concentration (MIC) and minimum biofilm eradication concentration (MBEC) values expressed in mg/mL. Values denoted as > “h.c.t.” indicate that MIC or MBEC exceeded the highest concentration tested. Representative assays for *C. cheilanthifolia* are shown below: **“plankton”** - planktonic growth, **“biofilm”** – biofilm growth, **SC** - sterility control, **GC** - growth control, solvent control - **DMSO** (bottom row).

The *in silico* toxicity assessment demonstrated that *Ch. majus* (both herb and root) showed the highest cumulative number of predicted toxic endpoints, indicating the broadest toxicological spectrum among the tested extracts. Moderate levels of predicted toxicity were observed for *F. officinalis* and *G. flavum* (herb and root), while *C. cheilanthifolia* (H1, H2, R) exhibited the lowest overall toxicity, with few or no predicted toxic endpoints across the evaluated categories.

Having established the basic pharmacokinetic and toxicological characteristics of the major alkaloids *in silico*, providing a preliminary but valuable insight into their potential applicability, we subsequently moved to experimental validation in the laboratory. As a first step, the antimicrobial activity of *Papaveraceae* extracts was assessed by determining their minimal inhibitory concentrations (MICs) and minimal biofilm eradication concentrations (MBEC) against reference ocular pathogens using the microdilution method in 96-well plates.

The tested plant extracts demonstrated variable activity against *S. aureus* and *P. aeruginosa*. In general, *S. aureus* was more susceptible, with MIC values ranging from 0.78 to 6.25 mg/mL and MBECs between 3.12 and 12.5 mg/mL. Extracts from *C. cheilanthifolia* (herb and root) and *G. flavum* root exhibited the strongest activity against *S. aureus*, with MICs of 0.78–1.56 mg/mL and MBECs of 3.12–12.5 mg/mL. In contrast, *P. aeruginosa* displayed reduced susceptibility, with several extracts (e.g., *Ch. majus* herb, *F. officinalis* herb, *G. flavum* herb and root) failing to achieve inhibition or eradication within the highest concentrations tested (h.c.t.).

Among the active samples, *C. cheilanthifolia* herb and *Ch. majus* root were the most effective, with MICs of 3.12–6.25 mg/mL and MBECs of 12.5 mg/mL; moreover Gram-positive *S. aureus* was generally more sensitive to the plant extracts than Gram-negative *P. aeruginosa*.

Building on these observations, we next employed a modified disc diffusion method (MDDM), in which bacterial cellulose (BNC) served as a carrier matrix for controlled release of the plant extracts. The resulting inhibition zones were irregular in shape; therefore, their surface areas (mm²) were quantified using ImageJ software instead of relying solely on diameter measurements (**Figure 5**, **Figure 6**).

**Figure 5.**
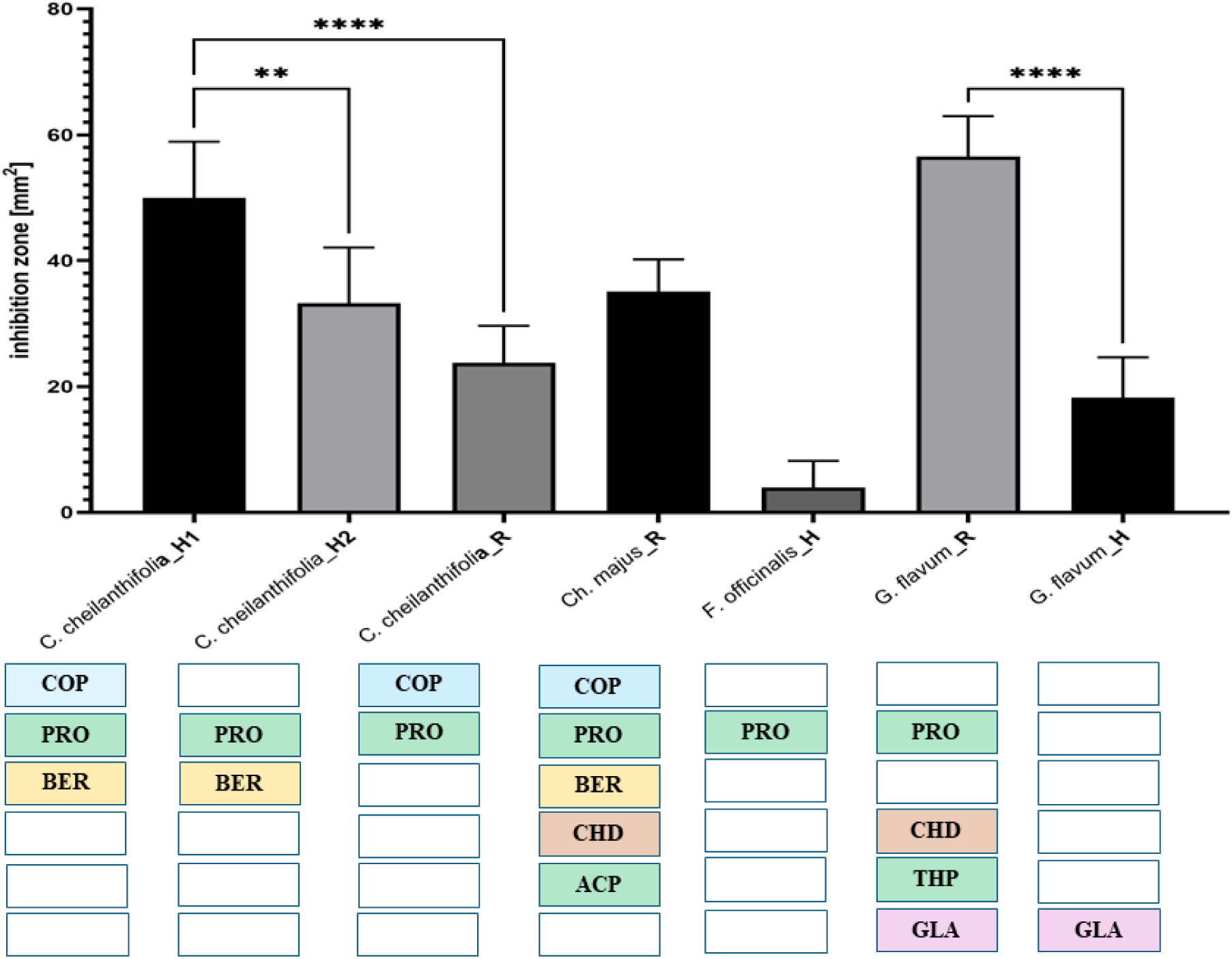
**Antimicrobial activity of ISQ alkaloid-rich extracts against *S. aureus.*** Inhibition zones (mm²) were quantified from six independent replicates using ImageJ software. The area corresponding to the 12-mm cellulose carrier was subtracted from each measurement to obtain the net inhibition zone. Data are expressed as mean ± SD (n = 6). Statistical significance was determined by one-way ANOVA followed by Tukey’s multiple comparison test (p < 0.01 – **, p < 0.0001 – ****; only selected comparisons are selected). The qualitative alkaloid composition of each extract, determined by LC–MS analysis, is indicated by abbreviations: **COP** – coptisine, **PRO** – protopine, **BER** – berberine, **CHEL** – chelidonine, **ACP** – allocryptopine, **THP** – tetrhydropalmatine, **GLA** – glaucine.

**Figure 6.**
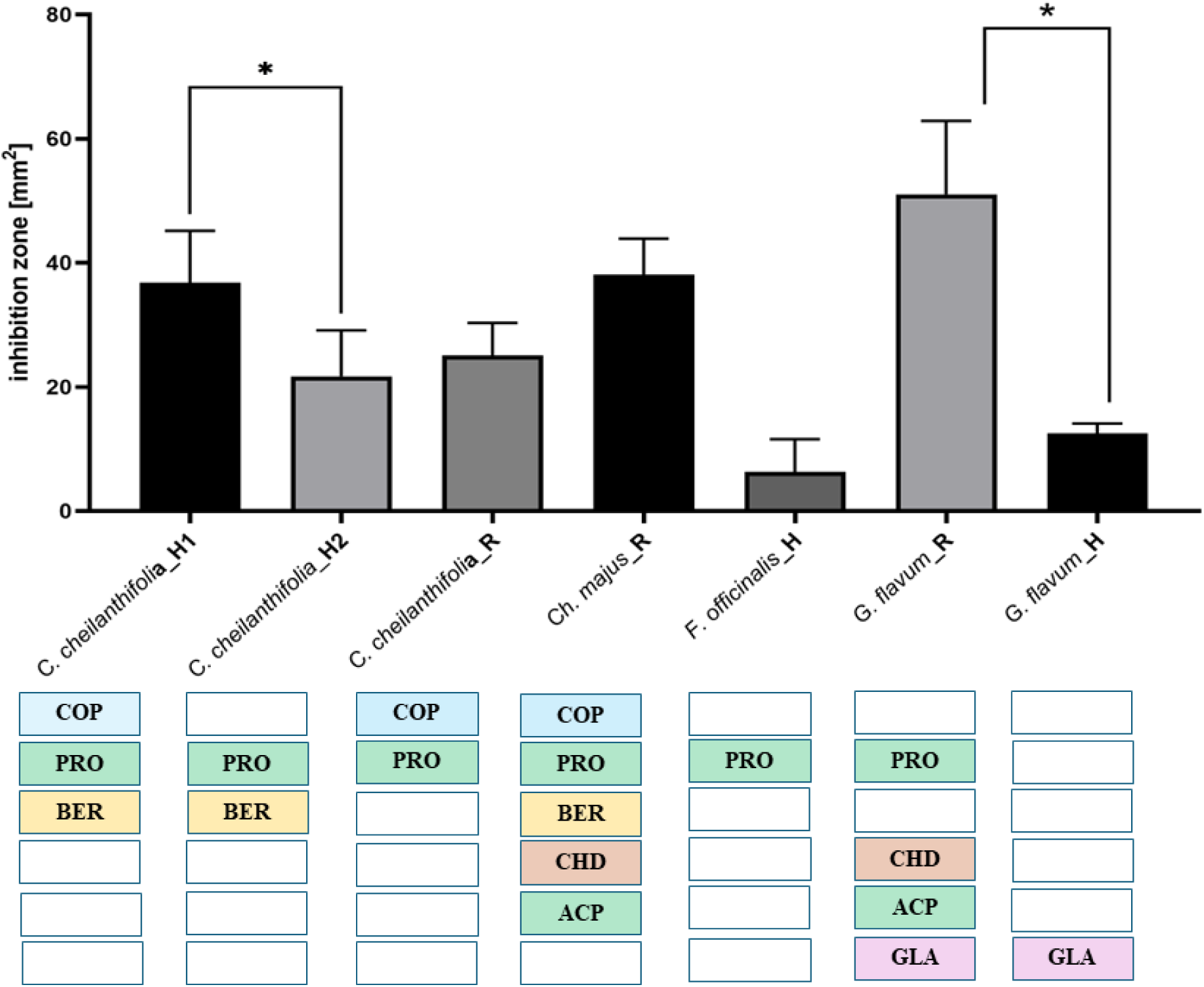
**Antimicrobial activity of ISQ alkaloid-rich extracts against *P. aeruginosa.*** Inhibition zones (mm²) were quantified from six independent replicates using ImageJ software. The area corresponding to the 12-mm cellulose carrier was subtracted from each measurement to obtain the net inhibition zone. Data are presented as mean ± SD (n = 6). Statistical significance was determined by one-way ANOVA followed by Tukey’s multiple comparison test (p < 0.05 – *, only selected comparisons are indicated). The qualitative alkaloid composition of each extract, determined by LC–MS analysis, is indicated by abbreviations: **COP** – coptisine, **PRO** – protopine, **BER** – berberine, **CHEL** – chelidonine, **ACP** – allocryptopine, **THP** – tetrhydropalmatine, **GLA** – glaucine.

The antimicrobial assay revealed distinct activity profiles of the tested *Papaveraceae* extracts against *S. aureus* and *P. aeruginosa*. Among all samples, *G. flavum_R* and *C. cheilanthifolia_H1* exhibited the largest inhibition zones for both bacterial species, indicating the strongest antibacterial potential. *C. cheilanthifolia_H2* and *C. cheilanthifolia_R* showed moderate activity, whereas *F. officinalis_H* produced only small inhibition zones. Overall, the results indicate that root extracts generally exhibited higher activity than herb extracts within the same species, and that *C. cheilanthifolia* and *G. flavum* extracts demonstrated the broadest and most potent antibacterial effects across both Gram-positive and Gram-negative strains.

Based on this data, BLISS and HAS analyses were performed to assess potential synergistic or additive, or antagonistic interactivities between particular alkaloids.

The interaction modeling based on Bliss independence and HSA criteria revealed distinct combinatorial behaviors of alkaloid mixtures across the tested Papaveraceae extracts. Against *S. aureus*, the most pronounced synergistic effects were observed for *C. cheilanthifolia_*H1 (COP + PRO + BER) and *G. flavum_R* (PRO + CHEL + THP + GLA), in which the observed inhibition zones exceeded Bliss predictions by more than 10%, indicating strong synergy between the constituent alkaloids. Moderate additivity was recorded for *C. cheilanthifolia_*H2 and *C. cheilanthifolia_*R, whereas *F. officinalis_*H, and *G. flavum_*H exhibited purely additive activity. The *Ch. majus_*R extract displayed partial antagonism in the Bliss model but weak synergy in HSA classification, reflecting divergence between models.

A comparable trend was noted against *P. aeruginosa*, where *C. cheilanthifolia_*H1 and *G. flavum_*R again showed the strongest synergistic interactions, while *C. cheilanthifolia_*H2 demonstrated moderate synergy under the HSA model and overall stable, reproducible antibacterial performance. The remaining extracts acted additively, with no antagonistic outcomes. Building on these interaction analyses, we next assessed the biocompatibility of the extracts *in vitro*. Since potential synergistic or additive effects between alkaloids could enhance not only antimicrobial efficacy but also cytotoxic potential, it was essential to evaluate their safety toward mammalian cells. Therefore, cytotoxicity testing was performed using established fibroblast cell models. The MTT assay was employed on fibroblasts as a first-line screening to determine the effects of individual and combined plant preparations on cell viability. This step provided an initial estimate of tolerable concentration ranges and helped distinguish selective antimicrobial activity from nonspecific cytotoxicity prior to further *in vivo* evaluation.

Cytotoxicity assay on L929 fibroblasts showed that most extracts maintained cell viability above the 80% threshold (horizontal line), indicating non-cytotoxic behavior according to norm ISO 10993-5. Only root extracts rich in protoberberine alkaloids caused a moderate reduction in viability. The solvent control (DMSO, ≤0.5% v/v) did not affect cell survival, confirming that the observed effects were due to the extracts themselves rather than the vehicle. Next, Bliss/HSA analysis was performed for cytotoxic assays *in vitro* [**Tab. 4**].

**Table 4.** Cytotoxic interaction modeling of alkaloid-containing plant extracts using Bliss independence and Highest Single Agent (HSA) reference models. Observed cytotoxic effects (expressed as 100 − cell viability) were compared to model-based predictions assuming a single global *E*ₘₐₓ. Interactions were classified as synergistic (SYN), additive (ADD), or antagonistic (ANT) within a ±10% tolerance range relative to the observed effect. All plant extracts showed low-to-moderate cytotoxicity toward fibroblasts, with predominantly additive or antagonistic profiles. Consistency between Bliss and HSA interpretations is indicated in the final column (Ag – agreement). COP - coptisine; PRO - protopine; BER - berberine; CHEL - chelidonine; ACP - allocryptopine; GLA- glaucine; THP - tetrahydropalmatine.

| Plant | Alk | Observed Cytotoxic Effect (%) | Bliss Prediction (%) | ΔBliss (Obs – Bliss) | Bliss Interpretation ($\pm 10\%$) | HSA Prediction (%) | ΔHSA (Obs – HSA) | HSA Interpretation ($\pm 10\%$) | HSA vs Bliss |
| --- | --- | --- | --- | --- | --- | --- | --- | --- | --- |
| *C. cheilant hifolia* H1 | cop +pro +ber | 27.65 | 30.55 | -2.9 | **ANT** | 30.55 | -2.9 | **ANT** | **Ag** |
| *C. cheilant hifolia* H2 | pro +ber | 30.14 | 30.55 | -0.4 | **ADD** | 30.55 | -0.4 | **ADD** | **Ag** |
| *C. cheilanthifolia* R | cop<br>+pro | 23.36 | 30.55 | -7.19 | ANT | 30.55 | -7.19 | ANT | Ag |
| *Ch. majus* R | cop<br>+pro<br>+ber<br>+chel<br>+acp | 21.95 | 30.55 | -8.6 | ANT | 30.55 | -8.6 | ANT | Ag |
| *F. officinalis* H | pro | 30.55 | 30.55 | 0.0 | ADD | 30.55 | 0.0 | ADD | Ag |
| *G. flavum* R | pro<br>+chel<br>+thp<br>+gla | 11.88 | 30.55 | -18.67 | ANT | 30.55 | -18.67 | ANT | Ag |
| *G. flavum* H | gla | 17.44 | 17.44 | -0.0 | ADD | 17.44 | -0.0 | ADD | Ag |

Bliss and HSA modeling, based on reconstructed single-alkaloid contributions, demonstrated that combinations containing (COP+BER) and (PRO+ACP) exhibited mainly additive or slightly synergistic cytotoxic effects. The pairing of PRO + CHEL or BER led to a moderate increase in observed cytotoxicity, exceeding Bliss predictions, which may indicate complementary mechanisms affecting cellular redox balance and membrane integrity. In contrast, interactions involving THPL and GLA in *G. flavum* extracts were antagonistic. Within *C. cheilanthifolia*, the comparison between H2 (PRO + BER) and H1 (COP + PRO + BER) extracts suggested that coptisine (COP) alone contributes modestly to cytotoxicity, yet enhances the overall effect in ternary mixtures - an example of weak synergy among protoberberine constituents. These findings indicate that the cytotoxic potential of compositions arises primarily from the additive accumulation of (COP, BER) and (PRO, ACP) activities, while the presence of (THP, GLA) alkaloids introduces antagonism, mitigating excessive toxicity. Given these results, *in vivo* validation was performed using the *G. mellonella* larval model to further assess systemic tolerance and potential toxicity under physiological conditions. This model provides a rapid and ethically compliant bridge between *in vitro* cytotoxicity and vertebrate safety testing, allowing integrated evaluation of both dose-dependent survival and visible tissue responses.

After injection, no mortality was observed in the control group of larvae (PBS + 1% DMSO) throughout 120 h. conversely, ethanol (75%) caused complete mortality (100%) within 24 h. Among the plant extracts, *F. officinalis*_H, *G. flavum*_H, and *C. cheilanthifolia*_H2 caused no mortality (0%) during the entire observation period, while *C. cheilanthifolia*_R and *G. flavum*_R induced lethality at the level of 16.6%. In turn, *C. cheilanthifolia*_H1 and *Ch. majus_R* caused higher lethality, reaching 33.2%. Next, before incorporating the extracts into the commercial formulation of artificial eye drops, their wettability was evaluated. Since the eye drops were intended to be subsequently embedded into NBC carriers, the primary property examined was their wettability, which essentially indicates how well the drop spreads across different surfaces (**Figure 8**).

**Figure 6.**
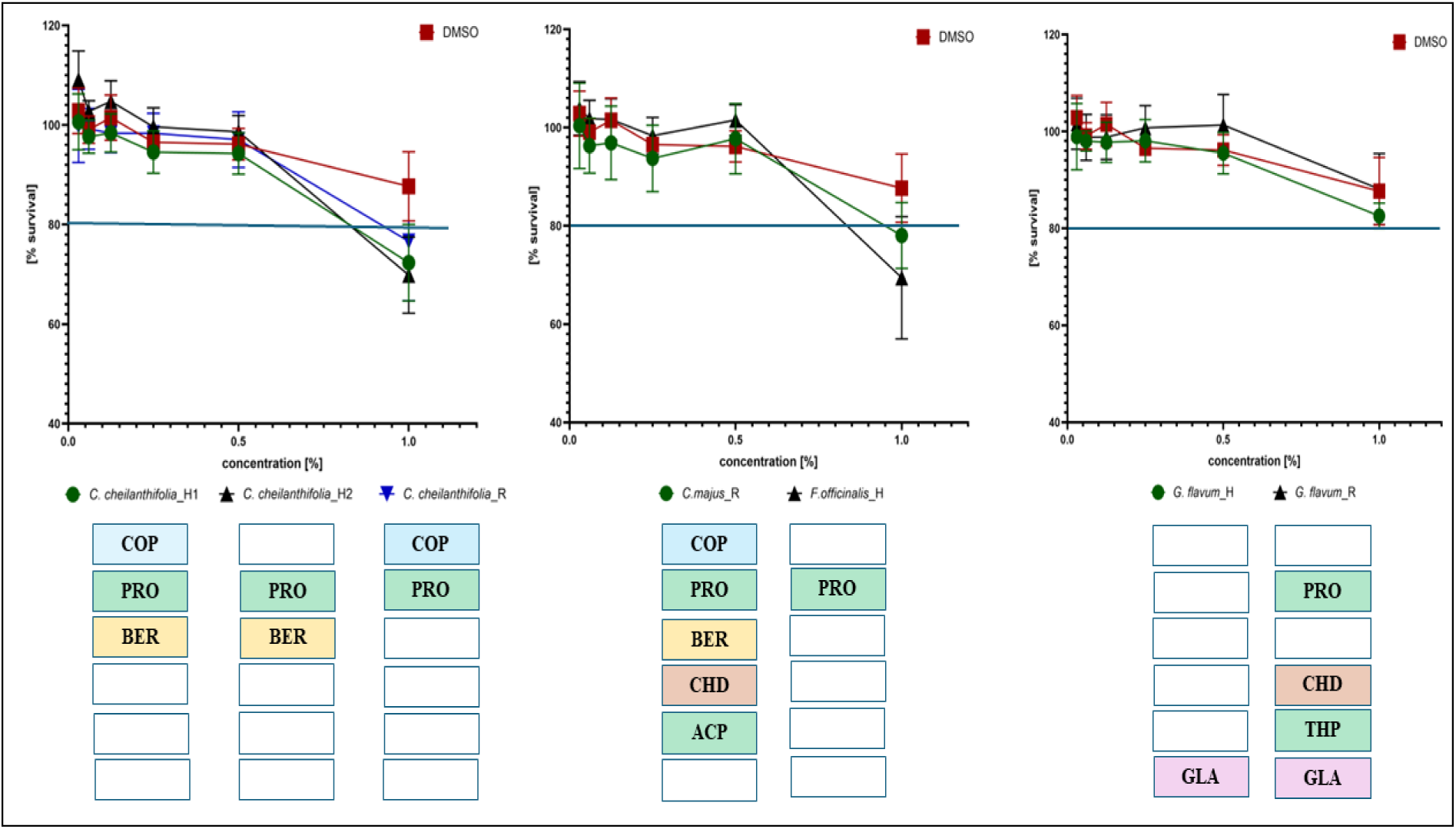
Cytotoxicity of plant preparations toward fibroblasts. Cells were exposed for 24 h to the indicated concentrations (0,03–1% v/v) of extracts; DMSO (red curve) is the matched solvent control at the same concentrations. Data are mean ± SD; the horizontal line marks 80% viability. Concentrations of 1% of the plant preparations corresponded to the concentration of 0.05mg/mL. Boxes under each plot list alkaloids detected in the corresponding extract: **COP**- coptisine; **PRO** - protopine; **BER** - berberine; **CHEL** - chelidonine; **ACP** - allocryptopine; **GLA** - glaucine; **THP** - tetrahydropalmatine.

**Fig. 7.**
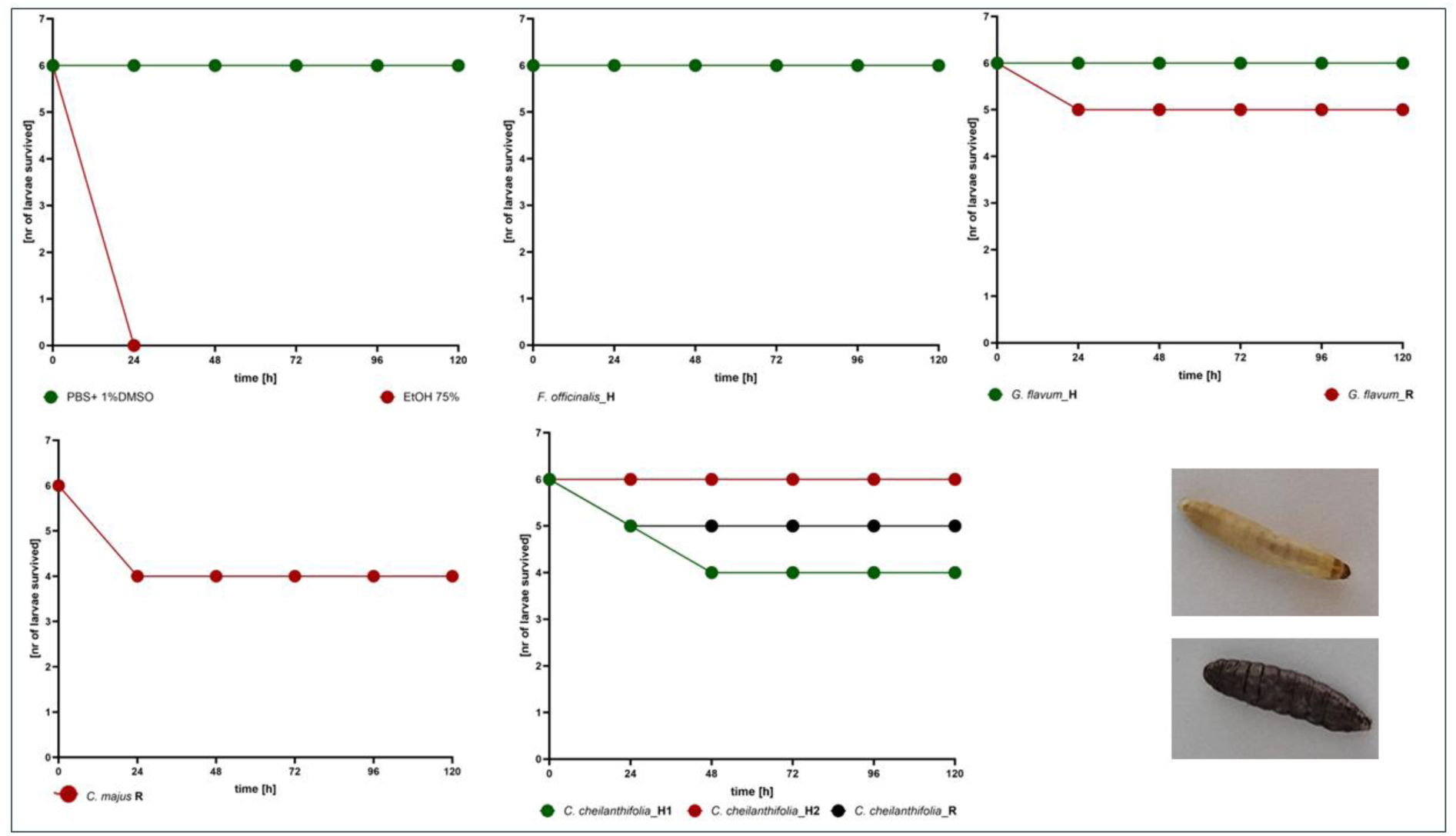
In v*i*vo toxicity assessment of plant extracts in the *G. mellonella* larval model. Larval survival was monitored for 120 h. PBS + 1% DMSO served as a negative control, while 75% ethanol acted as a positive cytotoxic reference. Representative images show a viable (upper image) and a melanized, non-viable (lower image) larvae.

**Fig. 8.**
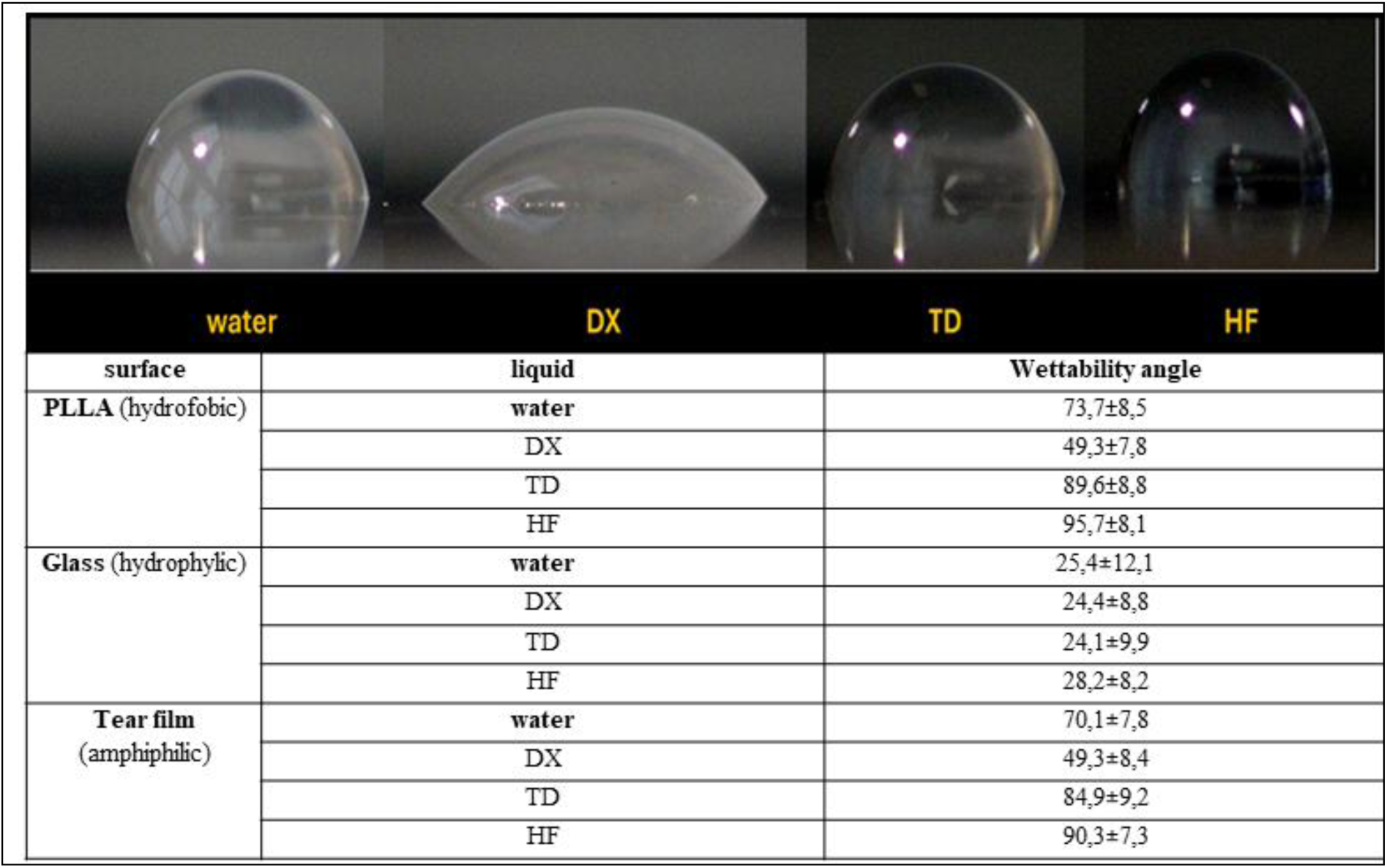
Wettability of **commercial ophthalmic formulations on surfaces with different physicochemical properties.** Representative images show sessile droplets DX, TD, and HF deposited on hydrophobic poly(L-lactic acid) (PLLA), hydrophilic glass, and an amphiphilic artificial tear film. Contact angles were determined using ImageJ. Lower contact angles correspond to higher wettability, indicating better spreading behavior of the tested eye drops on the given surface.

The results presented in **Fig.8** indicated that DX exhibited the most favorable spreading behavior, characterized by the lowest contact angles across all tested surfaces, suggesting optimal wettability and suitability for further incorporation into BC-based ocular carriers. Based on the integrated biological and physicochemical outcomes, the *C. cheilanthifolia_*H2 extract was selected for incorporation into the TX formulation. The resulting composite eye drops were subsequently embedded into BC membranes, and the outcomes of this stage are presented in **Fig.9**.

**Fig. 9.**
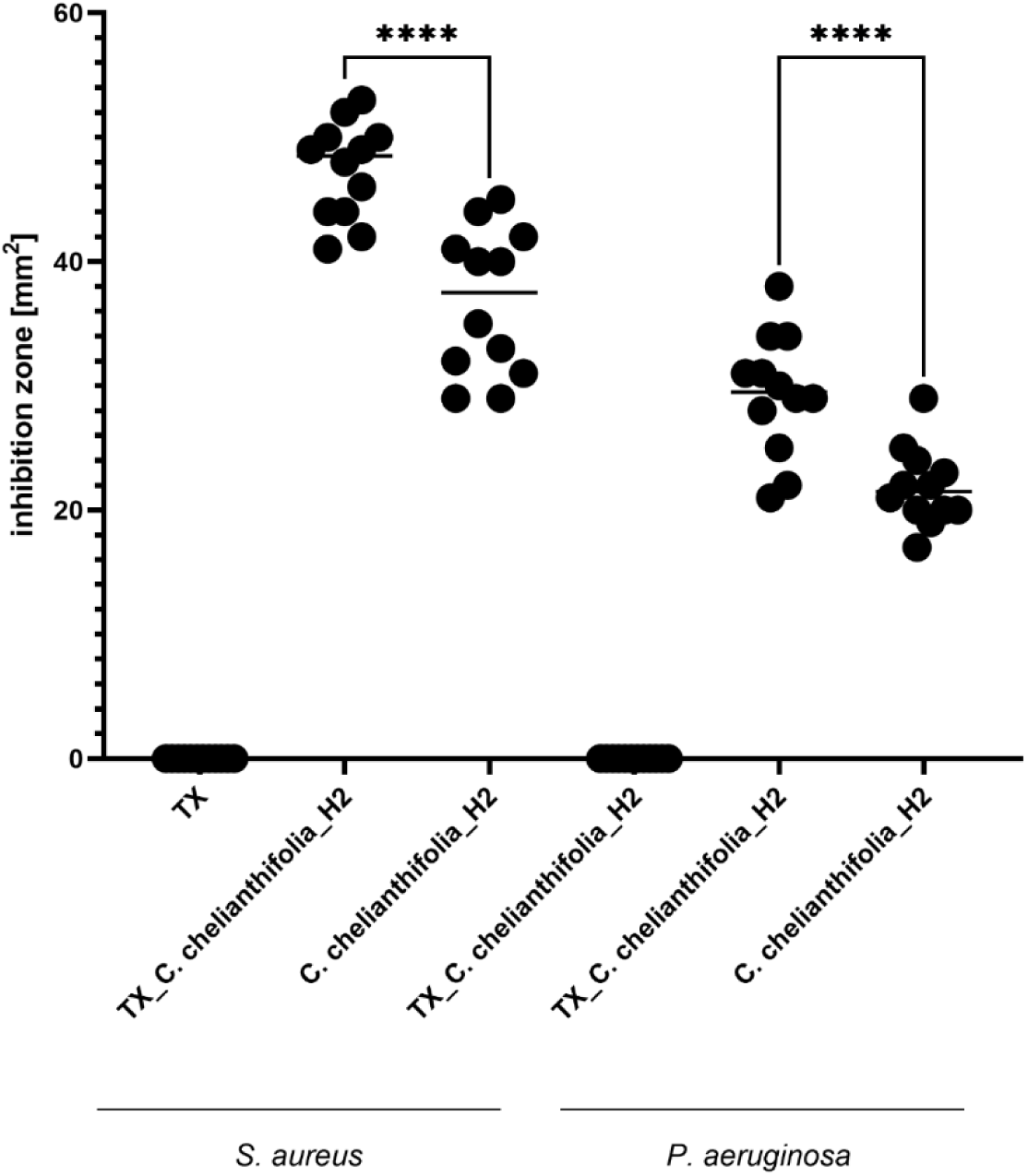
Antibacterial activity **of *Corydalis cheilanthifolia_H2* extract introduced to the TX eye drops containing compared to set of controls**. The modified disk diffusion assay was performed using NBC discs impregnated with TX + *C. cheilanthifolia_H2* or TD alone or *C. cheilanthifolia*_H2. Discs were placed on Mueller–Hinton agar plates inoculated with *S. aureus* ATCC 6538 or *P. aeruginosa* ATCC 15442. The inhibition zones were quantified in ImageJ, with the BC carrier area subtracted. Results represent the mean inhibition area ± SD from 12 repeats. Anova, followed with Sidak’s multiple comparison test; p<0.001.

The modified BC-based diffusion assay demonstrated that the antibacterial activity was driven by the *C. cheilanthifolia_H2* extract, while the TX vehicle containing 0.001% polyhexamethylene biguanide (PHMB) alone produced no inhibition zones against either *S. aureus* or *P. aeruginosa*. When the extract was incorporated into the TX formulation, the resulting inhibition zones were significantly larger than those generated by the extract alone (*p* < 0.05, Sidak’s multiple comparison test), for both tested strains. This finding indicates that the ophthalmic vehicle enhanced the bioavailability or diffusion of the active compounds without contributing intrinsic antimicrobial activity. The results confirm that the antibacterial potential of the *C. cheilanthifolia_H2* extract was not only preserved but further potentiated following its incorporation into the composite BC-based carrier.

## Discussion

The progressive loss of antibiotic efficacy due to bacterial resistance and the limited chemical diversity among available antiseptics justify the search for complementary antimicrobial strategies (Chinemerem Nwobodo et al., 2022; Muteeb et al., 2023). In our opinion, the studied plant-derived agents should not be regarded as substitutes for classical antimicrobials, but as a *third line of defense* - tools capable of temporarily relieving selective pressure while supporting infection control. The aim is not to replace, but to sustain and to provide locally active, short-contact compounds that reduce bacterial load without accelerating resistance. Within this framework, phytochemicals serve as a biological buffer zone between first-line (antibiotics) and second-line (antiseptics) interventions, extending the usability of both by preserving their clinical efficacy. In this work, the species from the Papaveraceae family were selected due to their unique and complementing phytochemical profiles, long ethnopharmacological history, and our group’s and others prior experience with its alkaloid constituents (Zielińska et al., 2021, 2019; Zhang et al., 2020, 2024; Bhambhani et al., 2021). These plants produce a diverse spectrum of isoquinoline alkaloids - structurally varied yet biosynthetically related - offering a unique model for studying additive, synergistic, and antagonistic interactions within natural mixtures (Krzyżek et al., 2021). *Ch. majus* is rich in benzophenanthridine, protoberberine, and protopine derivatives, particularly in its roots. Their concentrations may vary depending on the harvest time, habitat, and weather conditions. Numerous scientific reports indicate that the highest content and most diverse composition of isoquinoline alkaloids have been found in the roots of this species collected in autumn (Zielińska et al., 2019 and 2021; Tome and Colombo 1995). *C. cheilathifolia* as a species is relatively poorly described in the scientific literature. The few available studies indicate that, similarly to *Ch. majus*, it is a plant fairly rich in alkaloids, and its extracts exhibit strong antibacterial activity (Krzyżek et al., 2021; Zielińska et al., 2021). The richness of these compounds in both aerial and roots provides strong justification for including this species in studies aimed at developing a model set of plant metabolites with optimal antimicrobial activity. *G. flavum* and *F. officinalis* are characterized by a somewhat different alkaloid profile, which is associated with distinct biological properties (Wagner et al., 1996; Zhang et al., 2020; Akaberi et al., 2021; Kang et al., 2015). In both species, a high structural diversity of isoquinoline alkaloids has not been observed, unlike in *Ch. majus* or *C. cheilathifolia*, which makes them convenient model systems for comparative studies.

In turn, the ocular system was chosen as the testing ground because it imposes inherently strict pharmacodynamic conditions: continuous dilution (limited retention time) by tear fluid and exposure to opportunistic pathogens such as *S. aureus* and *P. aeruginosa*. Ocular infections remain a persistent clinical problem, particularly in postoperative and contact lens-related settings, where rapid microbial proliferation can compromise vision within dozen of hours (Shah and Wozniak, 2023). Therapeutic options in this context are restricted to a spectrum of antibiotics and antiseptics, both of which face aforementioned limitations related to tolerance, bioavailability, and resistance development. Thus, the clinical need is therefore not for stronger agents, but for safer, transiently active compounds that can reduce microbial burden without selecting for resistant strains or act together with antibiotics and antiseptics.

Nevertheless, it has to be noted that discourse surrounding local use of medical plants, including IQA-containing *Papaveraceae*, has long been polarized between two extremes: overestimated toxicity and exaggerated antimicrobial potency. Reports of hepatotoxicity, neurotoxicity, or mutagenicity are often generalized from extracts or isolated compounds administered systemically, without considering concentration, formulation, or route of exposure. Conversely, claims that these alkaloids kill Gram-negative and Gram-positive bacteria, as well as fungi, although true in general sense (such an activity is observed), often lack quantitative validation and also disregard critical variables such as concentration, exposure time or formulation, all of which profoundly influence the observed antimicrobial outcome. We ourselves - like many other research teams - have not entirely avoided this trap (Zielińska et al., 2020). This study sought to clarify these misconceptions by combining alkaloid profiling with biological analyses and to define the potential translational range of the extractible mixtures. Based on our previous studies, in which we used mixtures containing a limited number of components, namely two or three alkaloids in various combinations, to evaluate their antimicrobial effectiveness (Zielińska et al., 2019), we undertook investigations to develop a model mixture based on the correlation between the composition and the direction of bioactivity. Indeed, quantitative analysis of alkaloid content demonstrates that the total yield per 100 g of dry plant material was several orders of magnitude below doses associated with systemic toxicity in animal models (**Tab. 1**). Even under the conservative assumption of additive effects, the concentrations achievable in ocular formulations remain far below any toxic threshold. Moreover, given the small applied volume of eye drops (typically ≤ 5 mL) and the short exposure time before dilution (> 1minute, typically) and clearance by tear flow, the probability of reaching harmful tissue levels is negligible. Thus, while *Papaveraceae* alkaloids possess measurable bioactivity, their use in topically applied systems such as ophthalmic formulations should operate safely within a non-toxic window. *In silico* ADME modeling predicted that several major alkaloids, particularly berberine, chelidonine, allocryptopine, and protopine, may cross the blood–brain barrier and undergo hepatic metabolism (**Fig. 2**). However, again, these predictions must be interpreted in the context of the extremely low concentrations involved in topical ocular application. With dosing volumes of only a few milliliters and rapid dilution by tear turnover, systemic absorption is (if occurs at all) minimal. The possibility of local accumulation in ocular compartments such as the corneal epithelium or vitreous humor cannot be fully excluded (Lorrai et al., 2024), but given the rapid drainage kinetics of tear film, such effects would likely be transient.

Nevertheless, from a basic science perspective, the *in silico* toxicity modeling revealed an important relationship between alkaloid diversity within an extract and its predicted toxicological load: mixtures containing a greater number of distinct alkaloids exhibited higher cumulative toxicity scores (**Fig. 3**). The practical implication of this phenomenon once again indicates that while chemical complexity of extract increases theoretical toxicity, the administered doses and short exposure times in ophthalmic formulations should remain within the physiological safety margin.

Also, antimicrobial testing performed using both microdilution and modified disk diffusion assays (**Fig. 4–6**) revealed consistent activity patterns, showing that extracts containing more alkaloids generally exhibited stronger antibacterial effects. The microdilution method yielded quantitative MIC and MBEC values, providing a screening environment with efficient diffusion of the extract in liquid medium and prolonged contact with bacterial cells, thus ensuring reliable but initial detection of baseline antimicrobial activity. To address this limitation, a diffusion-based assay was employed, quantifying inhibition zones in mm². This method provided additional information, capturing not only antibacterial potency but also the compound’s ability to spread across soft and agar surfaces. When implemented with NBC as a carrier - which is also considered a potential dressing material for infected eyes (Malani et al., 2024) - the assay more accurately reproduced topical release conditions, enabling diffusion of alkaloids through the hydrated agar matrix. In our previous work, we have already demonstrated the favorable physicochemical and structural properties of a three-dimensional agar scaffold, particularly its capacity to mimic the characteristics of soft tissue (Ciecholewska-Juśko et al., 2022). It should be emphasized that modern, chemically synthesized antiseptics generally exhibit antimicrobial activity at much lower concentrations than these indicated for investigated here plant extracts (Kamaruzzaman et al., 2016; Papa et al., 2022; Franch et al., 2025). In this light, the activity of *Papaveraceae* extracts should be regarded as moderate but meaningful, especially considering their natural origin and multifactorial mechanisms of action.

In line with this, the formulation context plays a decisive role in shaping the apparent efficacy of *Papaveraceae* extracts. Interaction modeling using Bliss independence and Highest Single Agent (HSA) approaches demonstrated that *Papaveraceae* alkaloids exhibit a spectrum of interactions ranging from additive and synergistic (mostly) to antagonistic (rarely), depending on their specific composition. The strongest synergistic effects were observed in *C. cheilanthifolia_*H1 and *G. flavum_*R, where ternary or quaternary mixtures exceeded Bliss predictions by over 10%. Other extracts, such as *C. cheilanthifolia*_H2, acted mainly additively, offering a balanced compromise between activity and safety. The observed diversity in alkaloids’ composition allowed us, among others, to observe distinct patterns of synergism, additivity, and antagonism within naturally occurring alkaloid mixtures. These findings suggest that future (ours and of other teams) studies should adopt a reductionist strategy - systematically de-constructing the mixtures to identify which individual alkaloid contributes most to antimicrobial activity. Such knowledge would then enable a re-construction approach, where this principal compound could be formulated together with selected co-alkaloids providing cytoprotective or toxicity-buffering effects. This dual process of deconstruction and reconstruction may ultimately yield, (if combined with proper surfactants/emulgators) optimized, safer, and mechanistically transparent phytopharmaceutical formulations.

The ranking of antimicrobial potency among the tested extracts aligns with existing reports on the biological activity of individual *Papaveraceae* alkaloids. (Su et al., 2011; Zhang et al., 2023; Yang et al., 2025). The extracts richer in glaucine and tetrahydropalmatine, showed weaker or purely additive effects, mirroring literature data indicating their limited bacteriostatic capacity (Scholar, 2007; Du et al., 2022). The moderate but stable activity of *C. cheilanthifolia*_H2 fits within this framework, suggesting that extracts containing fewer, non-antagonistic alkaloids may achieve a clinically relevant antimicrobial profile.

I*n vitro* cytotoxicity assessment on L929 fibroblasts (**Fig. 6**) showed that most extracts maintained viability above the ISO 10993-5 safety threshold of 80%, confirming their overall biocompatibility. A moderate reduction in viability was observed only for *F. officinalis*_H, which was dominated by protopine. Although protopine has been reported by other teams to exert cytotoxic effects in certain cell types, in our opinion the observation presented in **Fig. 6** should be interpreted cautiously, as 2D cell culture systems are highly sensitive to mechanical detachment caused by viscous or surface-active solutions (Weiskirchen et al., 2023). The apparent loss of viability may therefore reflect partial cell detachment rather than genuine cytotoxicity. Importantly, this effect was absent in multi-alkaloid extracts such as *G. flavum*_R, suggesting also that the presence of other constituents may mitigate the response and support the overall safety of balanced plant extracts. Extracts characterized by lower phytochemical complexity, such as *F. officinalis*_H, showed higher cytotoxic indices, while those combining multiple structural classes - particularly *C. cheilanthifolia*_H2 and *G. flavum*_R - maintained cell survival within the non-cytotoxic range (**Tab. 4**). To verify whether these *in vitro* trends translate into a living organism, the *in vivo* toxicity test using *G. mellonella* larvae was subsequently performed and confirmed the overall safety of the evaluated extracts (**Fig. 7**). Complete survival (100%) was observed for *F. officinalis*_H, *G. flavum*_H, and *C. cheilanthifolia*_H2, while partial lethality (16.6–33.2%) occurred only for *Ch. majus*_R and *C. cheilanthifolia*_H1. It is noteworthy that these results do not fully parallel the *in vitro* data (**Fig.7** vs **Fig.6**) - fibroblast mortality was observed after exposure to *F. officinalis*, whereas no larval death occurred following injection of the same extract. This may further support our earlier assumption that cells in 2D cultures are more fragile and susceptible to mechanical or chemical stress than those embedded within an extracellular matrix or organized tissues of a living organism. Additionally, since several P*apaveraceae* alkaloids possess intrinsic anti-insecticidal properties (including activity against *G. mellonella*, specifically), the observed larval mortality (or its absence) may also reflect species-specific susceptibility rather than general cytotoxicity (Chelav and Khashaveh, 2013; Reyes-Luna et al., 2024). Nevertheless, the larval model, was in majority consistent with fibroblast cytotoxicity assays, and supported the favorable safety profile of *C. cheilanthifolia*_H2 as a translational candidate for ocular application. Integration of the selected extract into an ophthalmic formulation demonstrated that biological activity could be preserved and even enhanced under applied conditions. When *C. cheilanthifolia*_H2 was incorporated into the eye drop and further embedded within bacterial cellulose (BC) carriers, the resulting system exhibited superior spreading behavior and produced significantly larger inhibition zones against both *S. aureus* and *P. aeruginosa* compared to the extract or vehicle alone (**Fig 8–9**). The control formulation containing 0.001% PHMB showed no antibacterial effect, confirming that the observed activity originated from the plant extract itself. These findings validate preliminary the compatibility of *C. cheilanthifolia*_H2 with modern ophthalmic excipients and highlight its potential as a bioactive component of next-generation, surface-adherent therapeutic eye drops.

This study was designed as an exploratory translational framework, and several limitations should be acknowledged. Only two bacterial strains - *S. aureus* ATCC 6538 and *P. aeruginosa* ATCC 15442 - were examined. While they represent clinically relevant ocular pathogens, expanding the strain panel to include additional clinical isolates will be necessary to confirm broader applicability (Brożyna et al., 2025). The current results should therefore be interpreted as a conceptual reference point rather than a final antimicrobial profile. Moreover, although alkaloids were identified as the principal bioactive fraction, *Papaveraceae* extracts also contain flavonoids (quercetin, kaempferol, rutin, isorhamnetin) and phenolic acids (caffeic, ferulic, and *p*-coumaric acids), which may contribute to biological activity through antioxidant and membrane-modulating effects (Croft, 1998; Nazari et al., 2025). Generally, a higher content of polyphenolic compounds is characteristic of aerial parts, primarily the leaves, whereas underground plant organs, particularly in *Papaveraceae* species, contain substantially smaller amounts of these substances. In contrast, their alkaloid content is proportionally higher (Zielińska et al. 2021). Thus the focus on alkaloids was intentional, as they are the best-characterized pharmacophores of this family, allowing for clearer structure-activity interpretation. The cytotoxicity assay performed in this study relied on L929 fibroblasts, which, as a 2D epithelial-like model, may overestimate toxicity due to mechanical detachment of cells exposed to viscous or surface-active solutions. To mitigate this bias, complementary testing was conducted in *G. mellonella* larvae, though the insect model has its own limitations because certain *Papaveraceae* alkaloids possess intrinsic anti-insecticidal properties. Together, these approaches provide a conservative but informative view of safety under topical exposure conditions. The study also included a formulation containing polyhexanide (0.001%) as a preservative, yet a systematic analysis of synergy or additivity between the plant extract and antiseptics remains to be performed. Such work will be critical for designing combination therapies that enhance antimicrobial efficacy while minimizing the risk of resistance. Drug release from NBC was not examined, since the release kinetics of unmodified BC- rapid uptake and rapid release of small molecules -are well documented, including in our previous studies (Junka et al., 2019). Future stages will focus on implementing modified cellulose matrices, such as citric acid–crosslinked systems developed previously for wound applications, to achieve controlled release (Ciecholewska-Juśko et al., 2021). The physicochemical stability of the extracts within the TD formulation were also not assessed. These factors may influence diffusion and bioavailability and will be addressed in subsequent optimization studies. All these limitations are presented deliberately and explicitly, as each defines a clear direction that must be pursued before full translational implementation can be achieved. They also collectively illustrate why not all aspects could be addressed within a single study - no research team possesses the experimental capacity to comprehensively cover such a multidimensional framework at once.

Overall, this work constitutes a solid *in silico*, *in vitro*, and *in vivo* foundation that, once expanded to address the above limitations, will support targeted preclinical testing in vertebrate models and eventual translation into human ophthalmic applications.

## References

1. Akaberi, T., Shourgashti, K., Emami S.A., Akaberi, M. (2021). Phytochemistry and pharmacology of alkaloids from *Glaucium* spp. Phytochemistry, 191, 112923. 10.1016/j.phytochem.2021.112923

2. Aditi, S., Biharee, A., Kumar, A., Jaitak, V. (2020). Antimicrobial Terpenoids as a Potential Substitute in Overcoming Antimicrobial Resistance. Current Drug Targets, 21, 14. 10.2174/1389450121666200520103427

3. Ahmoda, R. A., Pirković, A., Milutinović, V., Milošević, M., Marinković, A., & Jovanović, A. A. (2025). Fumaria officinalis Dust as a Source of Bioactives for Potential Dermal Application: Optimization of Extraction Procedures, Phytochemical Profiling, and Effects Related to Skin Health Benefits. Plants, 14(3), 352. 10.3390/PLANTS14030352/S1

4. Akaberi, T., Shourgashti, K., Emami, S. A., & Akaberi, M. (2021). Phytochemistry and pharmacology of alkaloids from Glaucium spp. Phytochemistry, 191. 10.1016/J.PHYTOCHEM.2021.112923

5. Al-Doori, Z., Goroncy-Bermes, P., Gemmell, C. G., & Morrison, D. (n.d.). Low-level exposure of MRSA to octenidine dihydrochloride does not select for resistance. 10.1093/jac/dkm092

6. Anjali, Kumar, S., Korra, T., Thakur, R., Arutselvan, R., Kashyap, A. S., Nehela, Y., Chaplygin, V., Minkina, T., & Keswani, C. (2023). Role of plant secondary metabolites in defence and transcriptional regulation in response to biotic stress. Plant Stress, 8, 100154. 10.1016/J.STRESS.2023.100154

7. Bhambhani, S., Kondhare, K. R., & Giri, A. P. (2021). Diversity in Chemical Structures and Biological Properties of Plant Alkaloids. Molecules 2021, Vol. 26, Page 3374, 26(11), 3374. 10.3390/MOLECULES26113374

8. Brożyna, M., Stępnicka, Z., Kapczyńska, K., Dudek, B., Matkowski, A., & Junka, A. (2025). Establishing essential oil stewardship through the case of rosemary and thyme oils against Staphylococcus aureus. Frontiers in Microbiology, 16, 1668594. 10.3389/FMICB.2025.1668594/BIBTEX

9. Campano, C., Rivero-Buceta, V., Hernandez-Arriaga, A. M., Manoli, M. T., & Prieto, M. A. (2025). Pushing the limits of bacterial cellulose for biomedicine: a review. International Journal of Biological Macromolecules, 323, 146701. 10.1016/J.IJBIOMAC.2025.146701

10. Chakraborty, S., Chatterjee, R., & Chakravortty, D. (2022). Evolving and assembling to pierce through: Evolutionary and structural aspects of antimicrobial peptides. Computational and Structural Biotechnology Journal, 20, 2247. 10.1016/J.CSBJ.2022.05.002

11. Chelav, H. S., & Khashaveh, A. (2013). Insecticidal activity of Poppy (Papaver somniferum L.) seed oil against cowpea weevil (Callosobruchus maculatus F.) in stored cowpea. Archives of Phytopathology and Plant Protection, 46(19), 2314–2322. 10.1080/03235408.2013.792599

12. Chinemerem Nwobodo, D., Ugwu, M. C., Oliseloke Anie, C., Al-Ouqaili, M. T. S., Chinedu Ikem, J., Victor Chigozie, U., & Saki, M. (2022). Antibiotic resistance: The challenges and some emerging strategies for tackling a global menace. Journal of Clinical Laboratory Analysis, 36(9), e24655. 10.1002/JCLA.24655

13. Ciecholewska-Juśko, D., Żywicka, A., Junka, A., Drozd, R., Sobolewski, P., Migdał, P., Kowalska, U., Toporkiewicz, M., & Fijałkowski, K. (2021). Superabsorbent crosslinked bacterial cellulose biomaterials for chronic wound dressings. Carbohydrate Polymers, 253. 10.1016/J.CARBPOL.2020.117247

14. Ciecholewska-Juśko, D., Żywicka, A., Junka, A., Woroszyło, M., Wardach, M., Chodaczek, G., Szymczyk-Ziółkowska, P., Migdał, P., & Fijałkowski, K. (2022). The effects of rotating magnetic field and antiseptic on in vitro pathogenic biofilm and its milieu. Scientific Reports 2022 12:1, *12*(1), 1–19. 10.1038/s41598-022-12840-y

15. Cieplik, F., Jakubovics, N. S., Buchalla, W., Maisch, T., Hellwig, E., & Al-Ahmad, A. (2019). Resistance toward chlorhexidine in oral bacteria-is there cause for concern? Frontiers in Microbiology, 10(MAR), 431199. 10.3389/FMICB.2019.00587/TEXT

16. Croft, K. D. (1998). The chemistry and biological effects of flavonoids and phenolic acids. Annals of the New York Academy of Sciences, 854, 435–442. 10.1111/J.1749-6632.1998.TB09922.X

17. Deryabin, D., Galadzhiewa, A., Kosyan, D., Duskaev, G. (2019). Plant-Derived Inhibitors of AHL-Mediated Quorum Sensing in Bacteria: Modes of Action. Int. J. Mol. Sci. 2019, 20, 5588; doi:10.3390/ijms20225588

18. Du, Q., Meng, X., & Wang, S. (2022). A Comprehensive Review on the Chemical Properties, Plant Sources, Pharmacological Activities, Pharmacokinetic and Toxicological Characteristics of Tetrahydropalmatine. Frontiers in Pharmacology, 13, 890078. 10.3389/FPHAR.2022.890078/XML

19. Engman, S., Puello, F., Khoury, K., & Shah, D. (2023). S3801 Corydalis and Drug-Induced Liver Injury: A Series of Two Cases. American Journal of Gastroenterology, 118(10S), S2441–S2442. 10.14309/01.AJG.0000964844.89862.D1

20. Ezrari, S., Ben Khadda, Z., Boutagayout, A., Rehali, M., Jaadan, H., El Housni, Z., Khoulati, A., Saddari, A., & Maleb, A. (2025). Health risks and toxicity mechanisms of medicinal and aromatic plants (MAPs): A comprehensive review of adverse effects on organ systems, genotoxicity and reproductive toxicity. Fitoterapia, 184, 106630. 10.1016/J.FITOTE.2025.106630

21. Franch, A., Knutsson, K. A., Pedrotti, E., Fasolo, A., Bertuzzi, F., Birattari, F., Bonacci, E., Leon, P., & Papa, V. (2025). Treatment of Acanthamoeba keratitis with high dose PHMB (0.08%) monotherapy in clinical practice: A case series. European Journal of Ophthalmology, 35(4), 1235–1241. 10.1177/11206721241299470

22. Greater Celandine. (2022). LiverTox: Clinical and Research Information on Drug-Induced Liver Injury. https://www.ncbi.nlm.nih.gov/books/NBK548684/

23. Hardy, K., Sunnucks, K., Gil, H., Shabir, S., Trampari, E., Hawkey, P., & Webber, M. (2018). Increased usage of antiseptics is associated with reduced susceptibility in clinical isolates of Staphylococcus aureus. MBio, 9(3). 10.1128/MBIO.00894-18/ASSET/96CAC268-809F-40AD-AEDF-EF78FD9F273B/ASSETS/GRAPHIC/MBO0031839070004.JPEG

24. Hassan, K. A., Jackson, S. M., Penesyan, A., Patching, S. G., Tetu, S. G., Eijkelkamp, B. A., Brown, M. H., Henderson, P. J. F., & Paulsen, I. T. (2013). Transcriptomic and biochemical analyses identify a family of chlorhexidine efflux proteins. Proceedings of the National Academy of Sciences of the United States of America, 110(50), 20254–20259. 10.1073/PNAS.1317052110/SUPPL_FILE/PNAS.201317052SI.PDF

25. Junka, A., Żywicka, A., Chodaczek, G., Dziadas, M., Czajkowska, J., Duda-Madej, A., Bartoszewicz, M., Mikołajewicz, K., Krasowski, G., Szymczyk, P., & Fijałkowski, K. (2019). Potential of Biocellulose Carrier Impregnated with Essential Oils to Fight Against Biofilms Formed on Hydroxyapatite. Scientific Reports 2019 9:1, *9*(1), 1–13. 10.1038/s41598-018-37628-x

26. Kamaruzzaman, N. F., Firdessa, R., & Good, L. (2016). Bactericidal effects of polyhexamethylene biguanide against intracellular Staphylococcus aureus EMRSA-15 and USA 300. The Journal of Antimicrobial Chemotherapy, 71(5), 1252–1259. 10.1093/JAC/DKV474

27. Kang H., Jang S. W., Pak J. H., Shim S. (2015). Glaucine inhibits breast cancer cell migration and invasion by inhibiting MMP-9 gene expression through the suppression of NF-κB activation. Mol. Cell. Biochem, 403, 85–94. doi:10.3390/molecules27041378

28. Kim, M., Weigand, M. R., Oh, S., Hatt, J. K., Krishnan, R., Tezel, U., Pavlostathis, S. G., & Konstantinidis, K. T. (2018). Widely used benzalkonium chloride disinfectants can promote antibiotic resistance. Applied and Environmental Microbiology, 84(17). 10.1128/AEM.01201-18/SUPPL_FILE/ZAM017188706S1.PDF

29. Krzyżek, P., ……Zielińska, S. (2021). Helicobacter…

30. Li, X. L., Sun, Y. P., Wang, M., Wang, Z. Bin, & Kuang, H. X. (2024). Alkaloids in Chelidonium majus L: a review of its phytochemistry, pharmacology and toxicology. Frontiers in Pharmacology, 15, 1440979. 10.3389/FPHAR.2024.1440979

31. Lorrai, R., Cavaterra, D., Giammaria, S., Sbardella, D., Tundo, G. R., & Boccaccini, A. (2024). Eye Diseases: When the Solution Comes from Plant Alkaloids. Planta Medica, 90(06), 426–439. 10.1055/A-2283-2350

32. Malani, M., Thodikayil, A. T., Saha, S., & Nirmal, J. (2024). Carboxylated nanofibrillated cellulose empowers moxifloxacin to overcome Staphylococcus aureus biofilm in bacterial keratitis. Carbohydrate Polymers, 324. 10.1016/J.CARBPOL.2023.121558

33. Müller, G., & Kramer, A. (2008). Biocompatibility index of antiseptic agents by parallel assessment of antimicrobial activity and cellular cytotoxicity. Journal of Antimicrobial Chemotherapy, 61(6), 1281–1287. 10.1093/JAC/DKN125

34. Muteeb, G., Rehman, M. T., Shahwan, M., & Aatif, M. (2023). Origin of Antibiotics and Antibiotic Resistance, and Their Impacts on Drug Development: A Narrative Review. Pharmaceuticals, 16(11), 1615. 10.3390/PH16111615

35. Naghavi, M., Vollset, S. E., Ikuta, K. S., Swetschinski, L. R., Gray, A. P., Wool, E. E., Robles Aguilar, G., Mestrovic, T., Smith, G., Han, C., Hsu, R. L., Chalek, J., Araki, D. T., Chung, E., Raggi, C., Gershberg Hayoon, A., Davis Weaver, N., Lindstedt, P. A., Smith, A. E., … Murray, C. J. L. (2024). Global burden of bacterial antimicrobial resistance 1990–2021: a systematic analysis with forecasts to 2050. The Lancet, 404(10459), 1199–1226. 10.1016/S0140-6736(24)01867-1/ATTACHMENT/4F9FA746-A486-4BC0-BF71-075D4D1D6D56/MMC3.PDF

36. Nakamoto, M., Kunimura, K., Suzuki, J-I., Kodera, Y. (2020) Antimicrobial properties of hydrophobic compounds in garlic: *Allicin, vinyldithiin*, ajoene and diallyl polysulfides (Review) Experimental and Therapeutic Medicine 19: 1550-1553. DOI: 10.3892/etm.2019.8388

37. Nazari, N., Sharifnia, F., Salimpour, F., Mehrnia, M., Gran, A., & Zarafshar, M. (2025). Phytochemical richness, antioxidant activity, and molecular diversity in Papaver species from Iran’s western and central regions. Frontiers in Plant Science, 16, 1635867. 10.3389/FPLS.2025.1635867

38. O’Callaghan, R. J. (2018). The Pathogenesis of Staphylococcus aureus Eye Infections. Pathogens, 7(1), 9. 10.3390/PATHOGENS7010009

39. Papa, V., Van Der Meulen, I., Rottey, S., Sallet, G., Overweel, J., Asero, N., Minassian, D. C., & Dart, J. K. G. (2022). Safety and tolerability of topical polyhexamethylene biguanide: a randomised clinical trial in healthy adult volunteers. The British Journal of Ophthalmology, 106(2), 190–196. 10.1136/BJOPHTHALMOL-2020-317848

40. Reyes-Luna, A., Yáñez-Barrientos, E., Alba-Mares, X. N., Luis Olivares-Romero, J., Josabad Alonso-Castro, Á., Cruz Cruz, D., & Villegas Gómez, C. (2024). Metabolomic Approaches in Assessing the Insecticidal Activity of the Extracts from Argemone ochroleuca Sweet (Papaveraceae) Against Three Diverse Crop Pests of Economic Importance. Chemistry & Biodiversity, 21(2). 10.1002/CBDV.202301279

41. Saini, R.K., Ranjit, A., Sharma, K., Prasad, P., Shang, X., Gowda, K.G.M., Keum, Y.-S. (2022). Bioactive Compounds of Citrus Fruits: A Review of Composition and Health Benefits of Carotenoids, Flavonoids, Limonoids, and Terpenes. Antioxidants 2022, 11, 239. 10.3390/antiox11020239

42. Scholar, E. (2007). Glaucine. XPharm: The Comprehensive Pharmacology Reference, 1–4. 10.1016/B978-008055232-3.61825-2

43. Shah, S., & Wozniak, R. A. F. (2023). Staphylococcus aureus and Pseudomonas aeruginosa infectious keratitis: key bacterial mechanisms that mediate pathogenesis and emerging therapeutics. Frontiers in Cellular and Infection Microbiology, 13, 1250257. 10.3389/FCIMB.2023.1250257

44. Su, Y., Li, S., Li, N., Chen, L., Zhang, J., & Wang, J. (2011). Seven alkaloids and their antibacterial activity from Hypecoum erectum L. Journal of Medicinal Plants Research, 5(22), 5428–5432. http://www.academicjournals.org/JMPR

45. Teweldemedhin, M., Gebreyesus, H., Atsbaha, A. H., Asgedom, S. W., & Saravanan, M. (2017). Bacterial profile of ocular infections: a systematic review. BMC Ophthalmology, 17(1), 212. 10.1186/S12886-017-0612-2

46. Tome, F., and Colombo, M. L. (1995). Distribution of alkaloids in Chelidonium majus and factors affecting their accumulation. Phytochemistry 40, 37–39. doi: 10.1016/0031-9422(95)00055-C

47. Uruén, C., Chopo-Escuin, G., Tommassen, J., Mainar-Jaime, R. C., & Arenas, J. (2020). Biofilms as Promoters of Bacterial Antibiotic Resistance and Tolerance. Antibiotics, 10(1), 3. 10.3390/ANTIBIOTICS10010003

48. Wagner H, Bladt S. (1996). Plant Drug Analysis: A Thin Layer Chromatography Atlas. Second Edition. Springer-Verlag, Berlin, 9

49. Weiskirchen, S., Schröder, S. K., Buhl, E. M., & Weiskirchen, R. (2023). A Beginner’s Guide to Cell Culture: Practical Advice for Preventing Needless Problems. Cells 2023, Vol. 12, Page 682, 12(5), 682. 10.3390/CELLS12050682

50. Willcox, M. D. P. (2011). Review of resistance of ocular isolates of Pseudomonas aeruginosa and staphylococci from keratitis to ciprofloxacin, gentamicin and cephalosporins. Clinical & Experimental Optometry, 94(2), 161–168. 10.1111/J.1444-0938.2010.00536.X

51. Yang, X., Wang, Y., Li, L., Tang, D., Yan, Z., Li, M. Y., Jiang, J., & Bi, D. (2025). Berberine and its nanoformulations and extracts: potential strategies and future perspectives against multi-drug resistant bacterial infections. Frontiers in Microbiology, 16, 1643409. 10.3389/FMICB.2025.1643409/XML

52. Zhang, M., Yang, J., Sun, Y., & Kuang, H. (2024). Recent Advances in Alkaloids from Papaveraceae in China: Structural Characteristics and Pharmacological Effects. Molecules, 29(16), 3778. 10.3390/MOLECULES29163778/S1

53. Zhang, R., Guo, Q., Kennelly, E. J., Long, C., & Chai, X. (2020). Diverse alkaloids and biological activities of Fumaria (Papaveraceae): An ethnomedicinal group. Fitoterapia, 146. 10.1016/J.FITOTE.2020.104697

54. Zhang, R., Tian, S., Zhang, T., Zhang, W., Lu, Q., Hu, Q., Shao, H., Guo, Y., & Luo, Q. (2023). Antibacterial activity mechanism of coptisine against Pasteurella multocida. Frontiers in Cellular and Infection Microbiology, 13, 1207855. 10.3389/FCIMB.2023.1207855/BIBTEX

55. Zielińska S, Jezierska-Domaradzka A, Wójciak-Kosior M, Sowa I, Junka A and Matkowski AM (2018) Greater Celandine’s Ups and Downs−21 Centuries of Medicinal Uses of Chelidonium majus From the Viewpoint of Today’s Pharmacology. Front. Pharmacol. 9:299. 10.3389/fphar.2018.00299

56. Zielinska, S.; Wójciak-Kosior, M.; Dziagwa-Becker, M.; Glensk, M.; Sowa, I.; Fijałkowski, K.; Ruranska-Smutnicka, D.; Matkowski, A.; Junka, A. (2019). The Activity of Isoquinoline Alkaloids and Extracts From Chelidonium majus against Pathogenic Bacteria and *Candida* sp. Toxins, 11, 406.

57. Zielińska, S., Czerwińska, M.E., Dziągwa-Becker, M., Dryś, A., Kucharski, M., Jezierska-Domaradzka, A., Płachno, B., Matkowski, A. (2020). Modulatory effect of *Chelidonium majus* extract and its alkaloids on LPS-stimulated cytokine secretion in human neutrophils. Molecules, 25, 842; doi:10.3390/molecules25040842

58. Zielińska, S., Matkowski, A., Dydak, K., Czerwińska, M. E., Dziągwa-Becker, M., Kucharski, M., Wójciak, M., Sowa, I., Plińska, S., Fijałkowski, K., Ciecholewska-Juśko, D., Broda, M., Gorczyca, D., & Junka, A. (2021). Bacterial Nanocellulose Fortified with Antimicrobial and Anti-Inflammatory Natural Products from Chelidonium majus Plant Cell Cultures. *Materials (Basel*, Switzerland), 15(1). 10.3390/MA15010016

